# CXCL10 drives female-specific tau pathology progression and defines sex-dependent vulnerability in tauopathy model mice

**DOI:** 10.64898/2026.04.19.719088

**Authors:** Ryohei Uenishi, Rinna Kawata, Tatsuya Manabe, Yukio Matsuba, Naomi Mihira, Toru Takeo, Takaomi C. Saido, Masanori Hijioka, Takashi Saito

**Affiliations:** Department of Neurocognitive Science, Institute of Brain Science, Nagoya City University Graduate School of Medical Sciences, Nagoya, Japan; Department of Neuropathology, Graduate School of Medicine, The University of Tokyo, Tokyo, Japan; Laboratory for Proteolytic Neuroscience, RIKEN Center for Brain Science, Wako, Japan; Pioneering Research Division, Medical Innovation Research Center, Shiga University of Medical Science, Otsu, Japan; Division of Reproductive Engineering, Center for Animal Resources and Development, Kumamoto University, Kumamoto, Japan

**Author notes:** Corresponding authors: Correspondence to Drs. Masanori Hijioka and Takashi Saito. Contributions: R.U. designed and performed experiments, analyzed and interpreted data, drafted the manuscript, and acquired funding. R.K., Y.M., and N.M. performed experiments. T.M. analyzed and interpreted data, reviewed and edited the manuscript, and acquired funding. T.T. provided animal resources. T.C.S. provided animal resources, interpreted data, and reviewed the manuscript. M. H. designed and performed experiments, analyzed and interpreted data, reviewed and edited the manuscript, and acquired funding. T.S. conceived the study, designed experiments, analyzed and interpreted data, reviewed and edited the manuscript, acquired funding, and supervised the project. Ethics declaration: Competing interests: We declare no competing financial interests.

## Abstract

Neuroinflammation is a central driver of tauopathy, yet the precise chemokines that orchestrate the inflammatory microenvironment remain elusive. Here, we report C-X-C motif chemokine ligand 10 (CXCL10) is markedly upregulated in the brains of tauopathy model mice, where it co-localizes with prominent tau pathology. Notably, genetic ablation of *Cxcl10* in these mice significantly attenuates tau burden and extends the survival period, specifically in a female-dependent manner.

Mechanistically, although *Cxcl10* deficiency reduces the number of brain T cells in both sexes, this reduction does not correlate with the female-specific rescue of the phenotype. Furthermore, *Cxcl10* deficiency did not alter glial cell activation or motor function, suggesting a sex-specific mechanism. We show CXCL10 is primarily produced by pathological glia, fostering a localized inflammatory microenvironment. Our findings identify CXCL10 as a key mediator of tau pathology and reveal a sex-dimorphic regulatory axis that operates independently of T cell and glial activation paradigms.

## Introduction

Tauopathies are neurodegenerative disorders characterized by the abnormal accumulation of hyperphosphorylated and otherwise post-translationally modified tau protein within neurons and glial cells. These pathological tau species mislocalize and aggregate to form neurofibrillary tangles (NFTs), leading to distinct clinical phenotypes and disease classifications such as Alzheimer’s disease (AD), Pick’s disease (PiD), progressive supranuclear palsy (PSP), and corticobasal degeneration (CBD), depending on the affected tau isoforms and neuroanatomical regions ^1,2)^. While current therapeutic approaches, including tau-targeting antibodies and antisense oligonucleotides, are under intensive investigation, effective disease-modifying options remain elusive^3)^. The suboptimal outcomes of these tau-centric strategies underscore an urgent need to elucidate alternative or complementary mechanisms to address tauopathy.

Neuronal loss in tauopathies leads to macroscopic structural changes such as ventricular enlargement and cortical atrophy. Although NFT accumulation correlates with clinical progression, the mechanisms underlying neuronal death remain incompletely understood. Recently, the pathological role of glial cells and immune cells has emerged as a critical driver of disease progression and neurodegeneration. Glial subpopulations, including disease-associated astrocytes (DAA) and disease-associated microglia (DAM), have been identified in tauopathy model mice, and are thought to facilitate pathological progression^4–6)^. Furthermore, the increasing of immune cells, such as T cells, in the brain as a novel dimension of the pathophysiology of tauopathy. T cells are increased both in postmortem human brain tissue and murine models, and their depletion has been shown to mitigate tau pathology in model mice^7–10)^, suggesting that glial-immune interactions may serve as upstream modulators of disease progression.

Biological sex is another critical, yet poorly understood, variable in disease progression. Clinical observations indicate that adult females often exhibit faster progression of tau accumulation with cognitive decline in AD^11)^, whereas preclinical animal studies have reported sex-specific glial responses and neurodegeneration in tauopathy models, particularly in male mice^12)^. These disparities suggest that the molecular modulators of neuroinflammation may operate differently across sexes, necessitating sex-stratified investigations into glial-immune interactions.

Chemokines are essential regulators of immune cell migration and glial function. While altered chemokine levels have been reported in the cerebrospinal fluid (CSF) and brain tissues of patients with AD^13)^, the specific molecular mediators linking glial-derived chemokine signaling to T cell infiltration and tau pathology—especially in a sex-dependent manner—remain largely undefined. Clarifying these pathways may reveal novel, druggable targets that function independently of or synergistically with tau accumulation, while integrating sex-specific biological variables.

In this study, we identify and functionally characterize chemotactic factors driving disease progression in tauopathy model mice. Additionally, we examine how sex-specific differences contribute to immune cell dynamics and to the subsequent development of tau pathology. Through transcriptomic and biochemical analyses, we revealed a robust age-dependent upregulation of CXCL10 in the brains of P301S tau transgenic (Tau Tg) mice. CXCL10, a ligand for the chemokine receptor C-X-C motif chemokine receptor 3 (CXCR3), is a potent chemoattractant for T cells and macrophages^14,15)^ and is known to be upregulated in the CSF of patients with AD^16,17)^. Furthermore, CXCL10 has been reported to induce neurodegeneration by recruiting cytotoxic (CD8^+^) T cells in a multicellular neuroimmune culture system^18)^. These findings support our recent identification of CXCL10 as a candidate inflammatory factor contributing to the progression of tauopathy. We show here that CXCL10 is enriched specifically within brain regions susceptible to tau pathology. Notably, *Cxcl10* deficiency significantly reduced T cell accumulation in tauopathy models of both sexes. Critically, however, the suppression of tau pathology and the extension of lifespan upon *Cxcl10* deletion were observed exclusively in female mice, highlighting that CXCL10 plays a sex-dependent role in disease progression. These findings not only identify CXCL10 as a critical mediator of neuroimmune interaction in tauopathy but also implicate CXCL10 signaling as a driver of sex-based differences in disease severity. By uncovering this previously unrecognized immunopathogenic mechanism, our study provides a compelling rationale for targeting CXCL10 as a disease-modifying strategy, either alone or in combination with existing tau-directed therapies.

## Results

### CXCL10 is a predominant inflammatory mediator upregulated in the brains of tauopathy mice

To identify inflammatory mediators driving disease progression, we performed a comprehensive screening of the glial/immune landscape in Tau Tg mice, and found it to exhibit robust phosphorylated tau accumulation and progressive brain atrophy (Fig. 1a). Bulk RNA sequencing (RNA-seq) of the hippocampi from 12-month-old Tau Tg mice revealed a marked induction of inflammatory gene signatures (Fig.1b, c). Principal component analysis (PCA) and heatmap analysis confirmed distinct transcriptomic profiles in Tau Tg mice compared to wild-type (WT) controls, with gene set enrichment analysis (GSEA) highlighting the enrichment of diverse neuroinflammatory signaling pathways, led by the interferon (IFN) pathway (Fig. 1d, Fig. S1a, b). Notably, genes encoding key cytokines and chemokines were among the most significantly upregulated transcripts (Fig. 1e). To validate these findings at the protein level, we performed cytokine/chemokine arrays and enzyme linked immunosorbent assay (ELISA). We identified CXCL10 as a primary candidate as it showed a distinct increase in the brains of Tau Tg mice (Fig. 1f, g). Spatiotemporal analysis further demonstrated that CXCL10 protein levels were significantly elevated in the cortex and even more pronounced in the hippocampus as early as 9 months of age, persisting through 12 months in a sex-independent manner (Fig. 1h, i). Together, these data identify CXCL10 as a major inflammatory mediator associated with the progression of tau pathology.

**Fig. 1.**
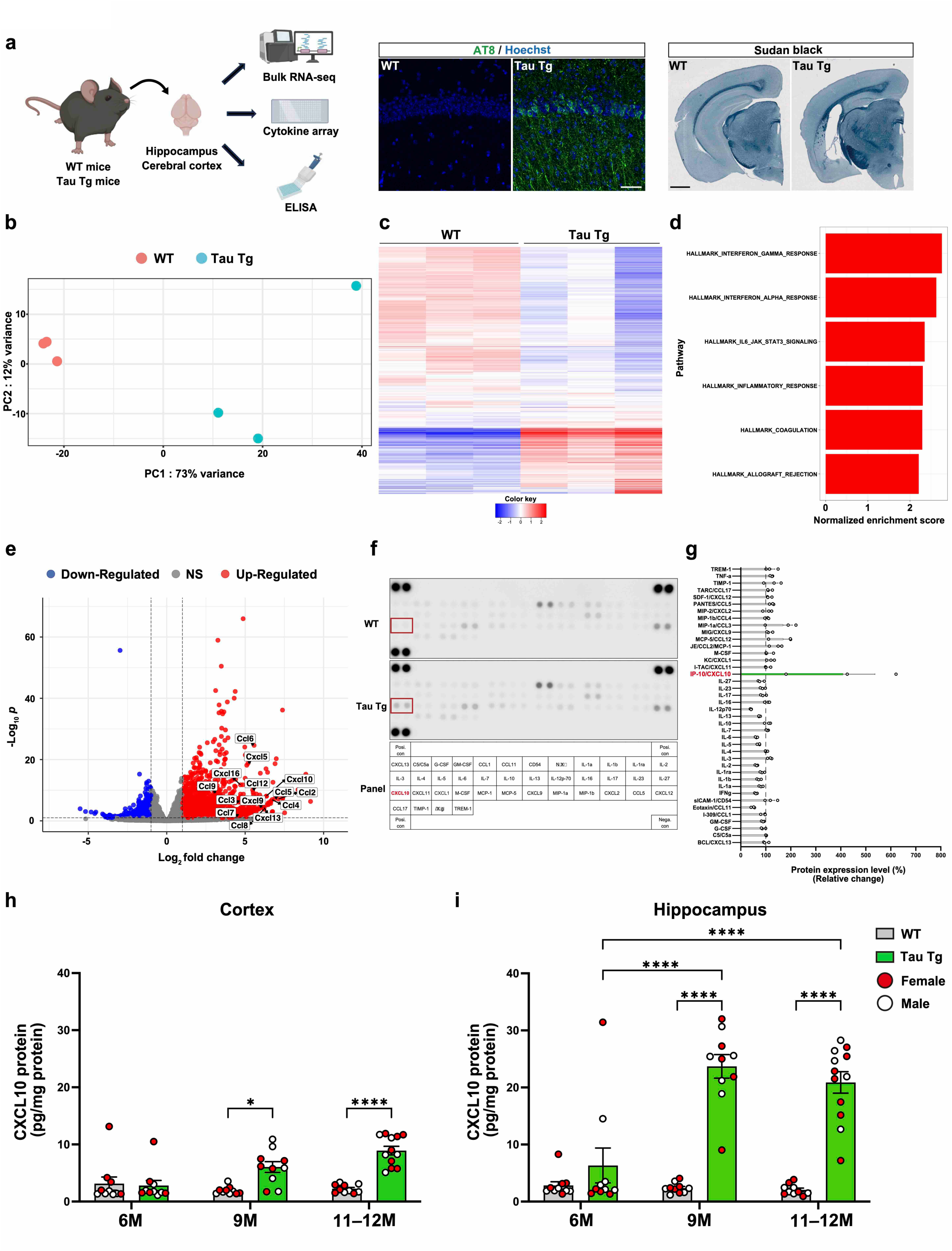
Gene and protein expression profiles of inflammatory molecules in the brains of Tau Tg mice. **A** Schematic of the experimental workflow. Hippocampi and cerebral cortices were collected from 12-month-old Tau Tg mice and age-matched WT controls. Bulk RNA-seq was performed on the hippocampi, while protein array analysis and CXCL10 ELISA were conducted using the cerebral cortices. At this age, Tau Tg mice exhibit prominent AT8-positive tau phosphorylation and brain atrophy. **b–e** Transcriptomic analysis of hippocampal bulk RNA-seq from Tau Tg and WT mice (n = 3 per group). **b** PCA plot. **c** Heatmap of the top 1,000 DEGs. **d** GSEA highlighting immune-related pathways enriched in Tau Tg mice. **e** Volcano plot of DEGs in Tau Tg mice compared to WT controls. DEG criteria: log₂ fold change > |1| and *p* < 0.1. **f, g** Cytokine and chemokine expression in the cerebral cortices of 9-month-old Tau Tg mice compared with age-matched WT controls. **f** Representative images of a cytokine/chemokine array using tissue lysates from the cerebral cortex. **g** Quantification of relative expression levels of indicated proteins in Tau Tg mice. Data are presented as mean ± S.E.M. **h, i** Expression levels of CXCL10 protein in the cerebral cortex (**h**) and the hippocampus (**i**) were measured by ELISA. Red dots indicate female mice, and white dots indicate male mice Data are presented as mean ± S.E.M. Number of mice used: except for 11–12-month-old female Tau Tg mice, where seven mice were used, we used five mice for each condition. For **b**–**e**, DEGs were identified using the Ward test. Statistical analysis was performed using a mixed-effects analysis followed by a Šídák’s multiple comparisons test (**h, i**). \**p* < 0.05, \*\*\*\**p* < 0.0001. Source data are provided in the Source Data file.

### *Cxcl10* deficiency transiently attenuates tau pathology and improves survival in a female-specific manner

Given the robust upregulation of CXCL10 during tau accumulation, we sought to determine its role in driving tau pathology. We crossed Tau Tg mice with *Cxcl10* KO mice and conducted sex-stratified analyses. At 9 months of age, both male and female Tau Tg mice showed substantial accumulation of pathological tau such as AT8-positive and sarkosyl-insoluble tau (Fig. 2a, f, g, l). While *Cxcl10* deficiency did not affect phosphorylated tau levels in males (Fig. 2a–f, m, n), female Tau Tg; *Cxcl10* KO mice exhibited a significant reduction in both AT8-positive and sarkosyl-insoluble tau, with other phosphorylated tau levels remaining unaltered (Fig. 2g–l, o, p). Notably, this protection in females translated to a survival benefit, as *Cxcl10* deficiency extended lifespan specifically in female mice (Fig. 2q, r). In males, while *Cxcl10* deficiency led to a transient extension of lifespan, these mice eventually succumbed to the pathology, reaching a mortality rate comparable to that of Tau Tg mice (Fig. 2q). Baseline tau pathology was largely comparable between sexes at this age, although males exhibited slightly higher total tau levels (Fig. S2a–c), suggesting that the suppressive effect of *Cxcl10* depletion on tau pathology is inherently a female-specific phenomenon.

**Fig. 2.**
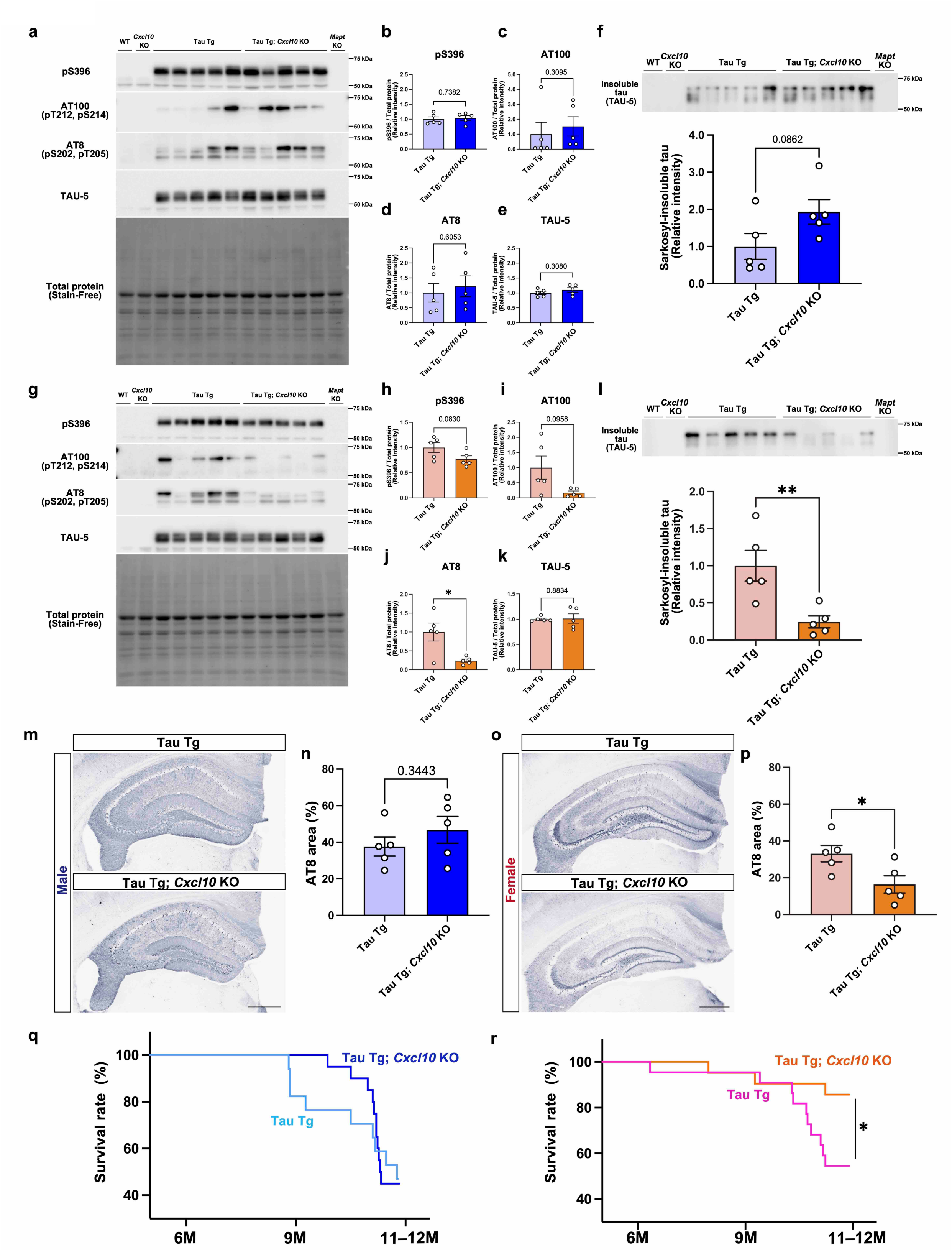
Female-specific modulation of tau pathology by *Cxcl10* deficiency in Tau Tg mice. a–l. Immunoblot analyses of tau protein in the cerebral cortices of 9-month-old Tau Tg mice with or without *Cxcl10* deficiency. Males (**a–f**), females (**g–l**). **a–e, g–k** Immunoblot images and relative expression of phosphorylated tau (p-tau) detected with anti-pSer396, pThr212/pSer214 [AT100], and pSer202/pThr205 [AT8] antibodies, and total tau detected with TAU-5 antibody. Protein expression was normalized to total protein using the Stain-Free method. **f, l** Immunoblot images of sarkosyl-insoluble tau from the cerebral cortex, including the piriform and entorhinal cortices. Bar graphs represent relative expression levels of tau protein in Tau Tg; *Cxcl10* KO mice compared to Tau Tg mice. **m**, **o** Immunohistochemical analysis of p-tau in the hippocampus using AT8 antibody. Bar graphs indicate the AT8-positive area in male (**n**) and female (**p**) mice. Scale bar = 500 µm. Data are presented as mean ± S.E.M. Number of mice used: Tau Tg (n = 5), and Tau Tg; *Cxcl10* KO (n = 5) for both sexes. **q**, **r** Graphs show survival curve in Tau Tg and Tau Tg; *Cxcl10* KO mice in males (**q**) and females (**r**). Number of mice used: Males: Tau Tg (n = 17) and Tau Tg; *Cxcl10* KO (n = 20); Females: Tau Tg (n = 22) and Tau Tg; *Cxcl10* KO (n = 21). Statistical analysis was performed using an unpaired two-tailed t-test (**b, d, e, f, h, l, n, p**), an unpaired t-test with Welch’s correction (**I, J, K**), a Mann-Whitney U test (**c**), and a log-rank test (**q**, **r**). \**p* < 0.05, \*\**p* < 0.01. Source data, including full blot images are provided in the Source Data file.

To determine the temporal window of this effect, we next examined 6-month-old mice with milder tau pathology (Fig. S3a, b). At this stage, AT8-positive tau accumulation was minimal and unaffected by *Cxcl10* deficiency in either sex (Fig. S4), likely reflecting the lack of significant CXCL10 induction at this early age (Fig. 1h, i). By 11–12 months, when tau pathology was markedly more advanced than at 9 months, the protective effect of *Cxcl10* deficiency in females was no longer observed (Fig. S3, S5). These findings suggest that CXCL10 exerts a transient, female-specific influence on the progression of tau pathology during a critical mid-stage window.

We also assessed whether CXCL10 modulates brain atrophy and neurodegeneration, given that significant brain atrophy has been reported in 11–12-month-old Tau Tg mice^19)^ Both male and female Tau Tg mice exhibited significant hippocampal atrophy and severe neuronal loss in the CA1, CA3, and DG regions; however, these phenotypes were not rescued by *Cxcl10* depletion (Fig. S6a–l). Given that tauopathy model mice have been reported to exhibit motor dysfunction alongside neurodegeneration^20)^, we evaluated functional deficits. Open field and rotarod tests detected no significant motor dysfunction and anxiety-like behavior in either sex (Fig. S7a–j). However, both sexes exhibited lower grip strength compared to WT mice, which was not rescued by *Cxcl10* deficiency (Fig. S7k–n). Additionally, severe hind limb abnormalities were observed in both male and female Tau Tg and Tau Tg; *Cxcl10* KO mice as evidenced by the foot clasping test (Fig. S7o-q). Thus, while CXCL10 transiently influences tau protein aggregation in females, its deficiency is insufficient to halt the terminal stages of neurodegeneration and motor decline in this model.

### Disease-associated astrocytes are the primary source of CXCL10 in tauopathy model mice

Disease-associated cell populations with distinct transcriptomic signatures preferentially localize to brain regions undergoing pathological progression in tauopathy model mice^21, 22)^. To accurately characterize these populations in a spatial context, we performed Xenium spatial transcriptomics using a 5,000-gene panel on the brains of 12-month-old WT and Tau Tg mice of both sexes (Fig. 3a). After quality control and data integration, uniform manifold approximation projection (UMAP) identified 23 distinct cell clusters (Fig. 3b). Based on publicly available single-cell RNA-seq (scRNA-seq) datasets^23)^, we annotated these clusters as neuronal cells, OPC-oligodendrocytes (Oligo), vascular cells (VC), border-associated macrophages (BAM), T cells (T), microglia (MG) and astrocyte-ependymal cells (AC-Epen). Intriguingly, we identified that *Cxcl10* mRNA is mainly expressed in AC-Epen and MG clusters (Fig. 3c). While per-cell expression levels were comparable, the number of *Cxcl10*-expressing cells was higher in Tau Tg brains than in WT controls in each cell type. (Fig. 3d). To achieve higher cellular resolution, we performed sub-clustering analysis defining the *Cxcl10*-expressing cell populations. AC-Epen were divided into 12 subcultures including eight AC subclusters as well as VC, Epen, and choroid plexus epithelial cells (CP) (Fig. 3e). Among these, AC8 emerged as a Tau Tg-specific cluster—regardless of sex—and was selectively distributed in the hippocampus and piriform cortex, these being regions exhibiting both prominent phosphorylated tau burden and extensive neurodegeneration (Fig. 3g, S8a, b). The AC8 transcriptome was markedly enriched with disease-associated astrocyte (DAA) markers, including *Serpina3n*, *C3*, and *Gfap* (Fig. S8c), and showed a distinct expression of *Cxcl10* in the Feature Plot (Fig. 3f).

**Fig. 3.**
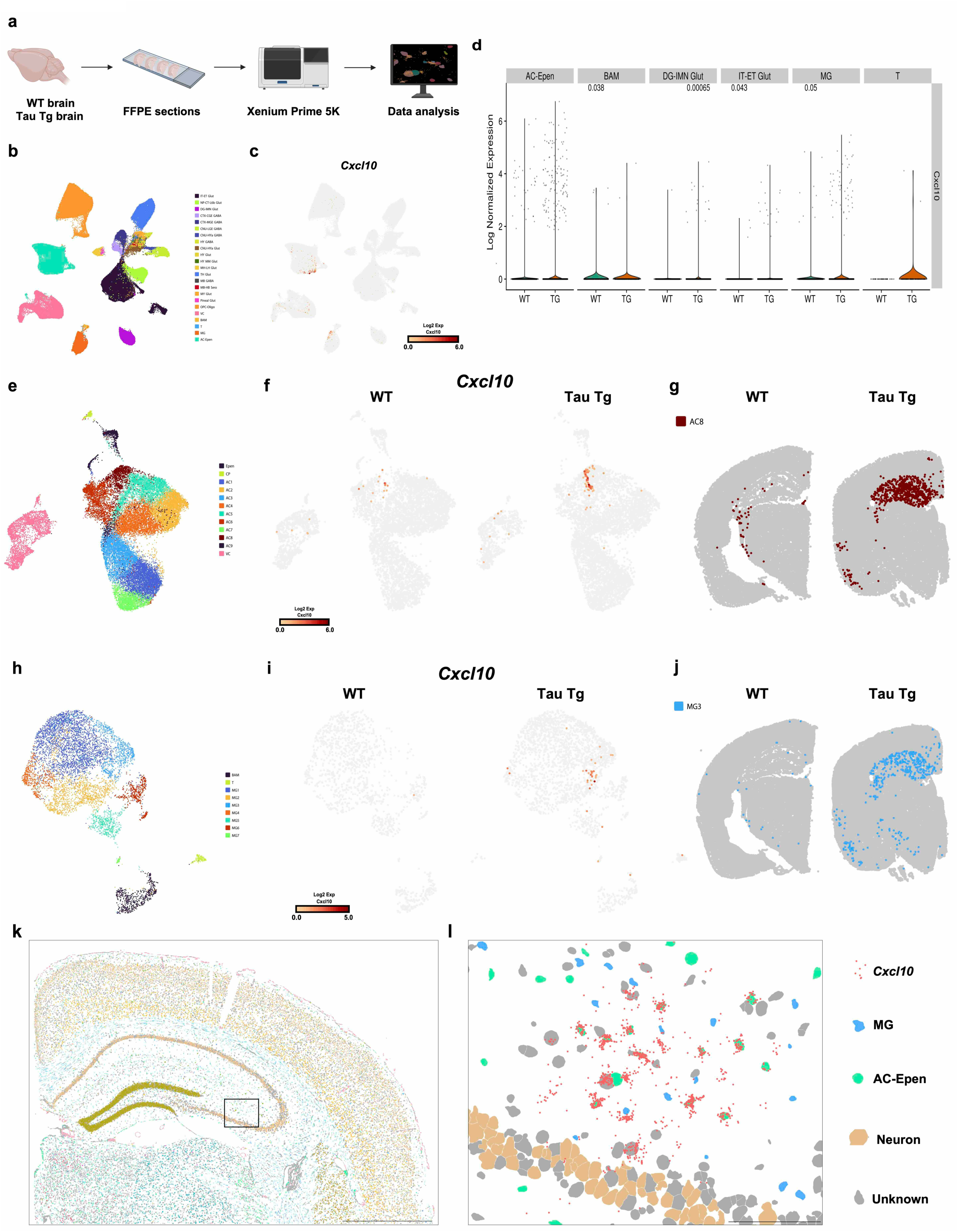
Pathological astrocytes are the main cell type expressing *Cxcl10* in Tau Tg mice. **a** Schematic diagram of the spatial transcriptome analysis using Xenium Prime 5K. Formalin fixed paraffin embedded (FFPE) sections were prepared from 12-month-old Tau Tg mice and age-matched WT controls. **b** UMAP visualizing the cell cluster detected by Xenium in the brains of Tau Tg and WT mice. Neuronal cells were classified as IT (intratelencephalic), ET (extratelencephalic), Glut (glutamatergic), NP (near-projecting), CT (corticothalamic), L6b (layer 6b), DG (dentate gyrus), IMN (immature neurons), CTX (cerebral cortex), CGE (caudal ganglionic eminence), GABA (GABAergic), MGE (medial ganglionic eminence), CNU (cerebral nuclei), LGE (lateral ganglionic eminence), Hya (anterior hypothalamic), HY (hypothalamus), MM (medial mammillary nucleus), LH (lateral habenula), TH (thalamus), MB (midbrain), HB (hindbrain), Sero (serotonergic), MY (medulla), NN (non-neuronal), NP (near-projecting), OB (olfactory bulb), OEC (olfactory ensheathing cells), and OLF (olfactory areas). **c** *Cxcl10* mRNA signal was plotted using Feature Plot on UMAP. **d** Quantitative *Cxcl10* gene expression using violin plots in AC-Epen, BAM, DG-IMN Glut, IT-ET Glut, MG, and T cell types. **e, h** Representative plots of the result of re-clustering AC-Epen (**e**) and immune cluster (**h**), respectively. **f** Plots of *Cxcl10*^+^ cells in the cluster shown in Fig. 3e and 3h represented according to genotype. **g, j** Figures showing spatial distribution of AC8 (**g**) and MG3 (**j**) clusters in the brains of WT and Tau Tg mice. **k** Representative images of coronal section of mouse brain by Xenium explorer. Scale bar = 1 mm. **l** Spatial information of *Cxcl10*^+^ astrocytes and microglia in the hippocampus of Tau Tg mice using Xenium explorer. Scale bar = 100 μm. Number of mice used: male WT (n = 1), male Tau Tg (n = 1), female WT (n = 1), and female Tau Tg (n = 1). Statistical analysis was performed using a Wilcoxon rank sum U statistic test (**d**). Source data are provided in the Source Data file.

Similarly, we also identified immune cell clusters (MG, T, BAM) were divided into 9 subclusters, including seven MG subclusters, T, and BAM (Fig. 3h). Among these, the MG3 emerged as a distinct cluster in Tau Tg mice, which localized to similar regions as those with phosphorylated tau (Fig.3 j, S9a, b). The MG3 expressed *Cxcl10* mRNA along with disease-associated microglia (DAM) signatures, such as *Cst7*, *Axl*, and *Itgax* (Fig. S9c). Notably, we observed that the number of *Cxcl10*^+^ astrocytes significantly outweighed that of *Cxcl10*^+^ microglia within the CXCL10-rich niche (Fig. 3k, l). Therefore, *Cxcl10* is mainly expressed in pathological astrocytes characterized by DAA-associated genes and selectively localized to brain regions with prominent tau pathology. Furthermore, expression of *Cxcr3*, the receptor for CXCL10, was identified primarily in MG, BAM, and most prominently in T cells (Fig. S10a, b).

To validate these spatial findings, we conducted RNAscope in combination with immunohistochemical analysis (IHC). In the hippocampus and piriform/entorhinal cortex, *Cxcl10* mRNA signals were highly abundant in Tau Tg mice of both sexes (Fig. S11a, b). Co-staining confirmed that the vast majority of *Cxcl10* mRNA colocalized with GFAP^+^ astrocytes, with only sparse signals in Iba1^+^ microglia, while *Cxcr3* mRNA was mainly detected in CD3^+^ T cells (Fig. S11c, d). These findings were further corroborated by re-analysis of public single-nucleus RNA-seq (snRNA-seq) data from PS19 mice (GSE221856) ^24)^, which showed significant *Cxcl10* upregulation in the astrocyte cluster but not in the microglia cluster (Fig. S12a–e). Next, re-analysis of the GSE218728 dataset^8)^ confirmed *Cxcr3*-expressing cell types in the brain. Consistent with our observations, *Cxcr3* mRNA was identified in T and NK cells, but not in microglia, B cells, or neutrophils (Fig. S13a–d). In addition, the proportion of T cells in the brain was higher than that of NK cells (Fig. S13e). Together, these findings demonstrate that pathological astrocytes are the major cellular source of CXCL10, whereas CXCR3 is primarily expressed in T cells in the tauopathy model mice.

### *Cxcl10*^+^ astrocytes activate inflammatory responses in the surrounding microenvironment in the brains of tauopathy model mice

While CXCL10 was predominantly expressed in pathological astrocytes in the brains of Tau Tg mice, its expression was heterogeneous across the astrocyte population. To elucidate the functional impact of astrocyte-derived CXCL10 on the surrounding microenvironment for associating tau pathology formation, we compared the transcriptomic profiles of cells located within a 50 µm radius of either *Cxcl10*^+^ or *Cxcl10*^-^astrocytes (Fig. 4a). Spatial vicinity analysis found that T cell marker genes, including *Cd3d* and *Cd3g*, were upregulated in the vicinity of *Cxcl10*^+^ astrocytes compared to their *Cxcl10*^-^ counterparts. This enrichment was observed in male Tau Tg mice and followed a consistent trend in females (Fig. 4b, S14a). Next, we performed pathway analysis to elucidate functional differences in these cells. Gene Ontology (GO) analysis showed an enrichment for terms related to responses to cytokines and chemokine receptor binding in both sexes (Fig. 4c, d, S14b, c). Furthermore, we revealed that the inflammatory response is more activated in both sexes in the microenvironment where CXCL10 is present (Fig. 4c, S14b). These results suggest that *Cxcl10*^+^ astrocytes actively recruit or interact with T cells likely expressing *Cxcr3* in the hippocampus in both sexes. Furthermore, CXCL10 may serve as an important regulator of the inflammatory microenvironment within the tauopathy mouse brain.

**Fig. 4.**
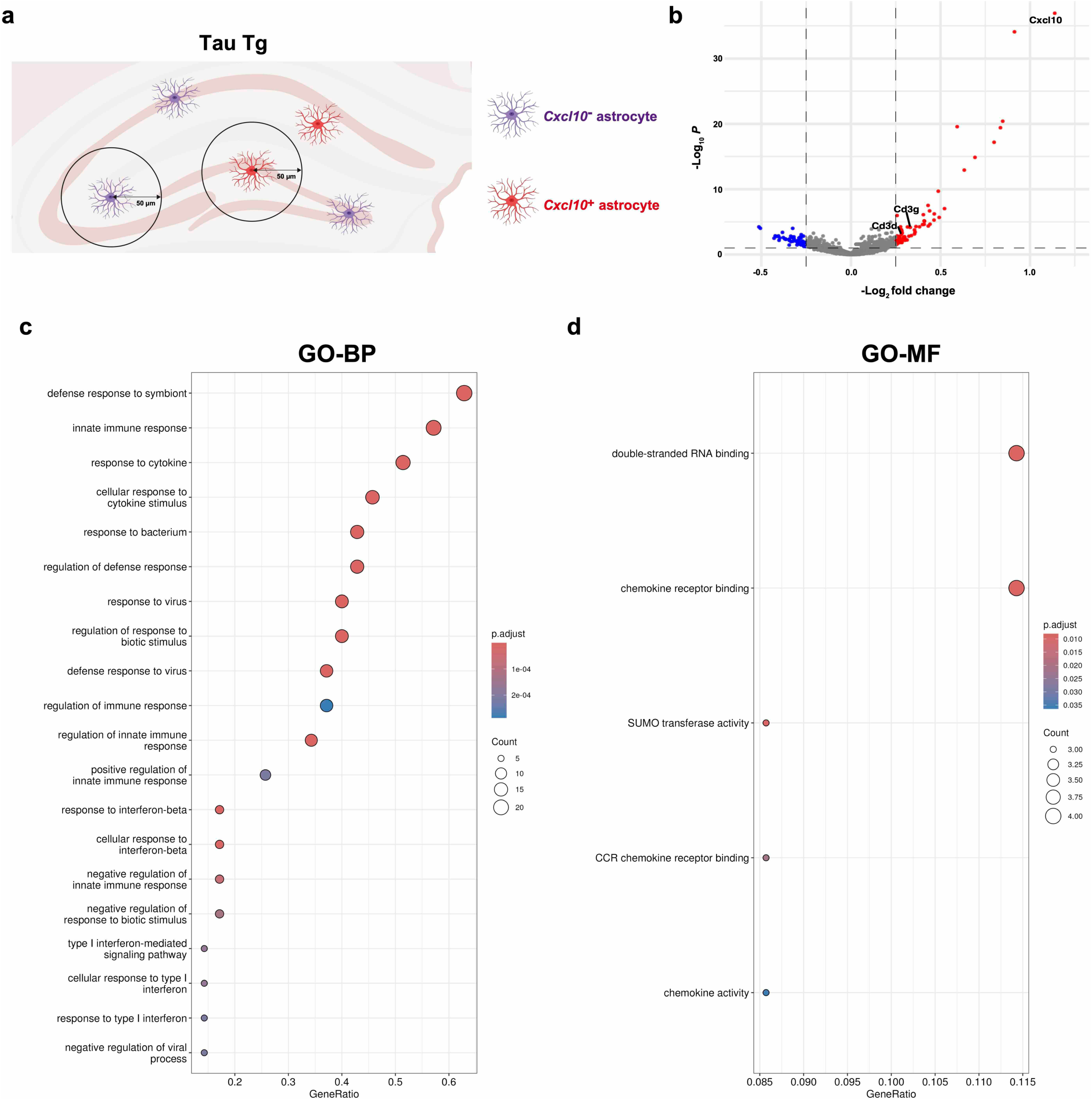
The local microenvironment where CXCL10 is present activates inflammatory response pathways in tauopathy model mice. **a** Illustration explaining the definitions of cells neighboring *Cxcl10*^+^ astrocytes and *Cxcl10*^-^ astrocytes. **b** Volcano plot of gene expression showing Log₂ fold change > |0.25| and p < 0.1 highlighted in red, and Log₂ fold change < |0.25| and p > 0.1 shown in blue. All other transcripts are indicated in gray. **c** GO pathway analysis focusing on biological processes (BP) and molecular functions (MF) performed to compare neighboring cells of *Cxcl10*^+^ astrocytes with neighboring cells of *Cxcl10*^-^ astrocytes in male Tau Tg mice. The top 20 significantly upregulated pathways in neighboring cells of *Cxcl10*^+^ astrocytes are shown in each panel. Number of mice used: male Tau Tg (n = 1). (**b**–**d**) DEGs were identified using the Wilcoxon test, and GO analysis was evaluated via over-representation analysis employing the hypergeometric test. Source data are provided in the Source Data file.

### CXCL10 partially affects brain T cells but not glial activation in Tau Tg mice

Since immune-related genes were upregulated in cells adjacent to *Cxcl10*^+^ astrocytes (Fig 4a–c), we next investigated whether CXCL10 modulates immune cell dynamics, specifically T cells, in Tau Tg mice. First, we performed flow cytometry (FCM) to analyze the populations of CD3^+^, CD4^+^, and CD8^+^ T cells in the brains of Tau Tg and Tau Tg; *Cxcl10* KO mice (Fig. S15a, b). We found that one population of brain T cells, specifically CD8^+^ T cells, increased in Tau Tg mice at 11–12 months of age in both sexes. However, *Cxcl10* depletion did not significantly alter the overall T cell populations compared to Tau Tg mice (Fig. S15c–h). To examine immune cell dynamics in regions with marked tau pathology, we also performed IHC specifically on the hippocampus. We found a dramatic increase in CD3^+^ T cells from 9 to 11–12 months in both sexes. In contrast, *Cxcl10* deficiency partially reduced the number of T cells in 11–12-month-old mice of both sexes (Fig. 5a–d). Notably, T cells did not increase in the brains of 9-month-old female Tau Tg mice, a stage where protective effects are already emerging. These data suggest that the neuroprotective effects of *Cxcl10* deficiency on tau accumulation and survival in females are unlikely to be primarily mediated by the modulation of T-cell populations. To further explore how *Cxcl10* depletion affects tau pathology, we examined microglia and astrocytes, both of which are key contributors to the progression of tauopathy^25)^. We quantified areas within the CA1 and DG of the hippocampus occupied by Iba1^+^ microglia and complement C3^+^ astrocytes, the latter of which is known to be highly expressed in reactive astrocytes in PS19 mice^26)^. Microglial and astrocytic activation was initiated at 9 months in male Tau Tg mice and became prominent in both sexes by 11–12 months. However, *Cxcl10* deficiency did not significantly alter glial activation compared to age-matched Tau Tg mice (Fig. S16a–d). In addition, border-associated macrophages (BAMs) expressing *Cxcr3* mRNA showed no apparent changes in cell number or expression levels in the hippocampus and piriform cortex of Tau Tg mice, suggesting that BAMs do not contribute to tau pathology progression (Fig. S10b, S17). Collectively, these results indicate that neither glial cells nor BAMs are primarily involved in the protective effects observed in female Tau Tg; *Cxcl10* KO mice. This suggests that the observed benefits of *Cxcl10* deficiency may be modulated by other unidentified factors.

**Fig. 5.**
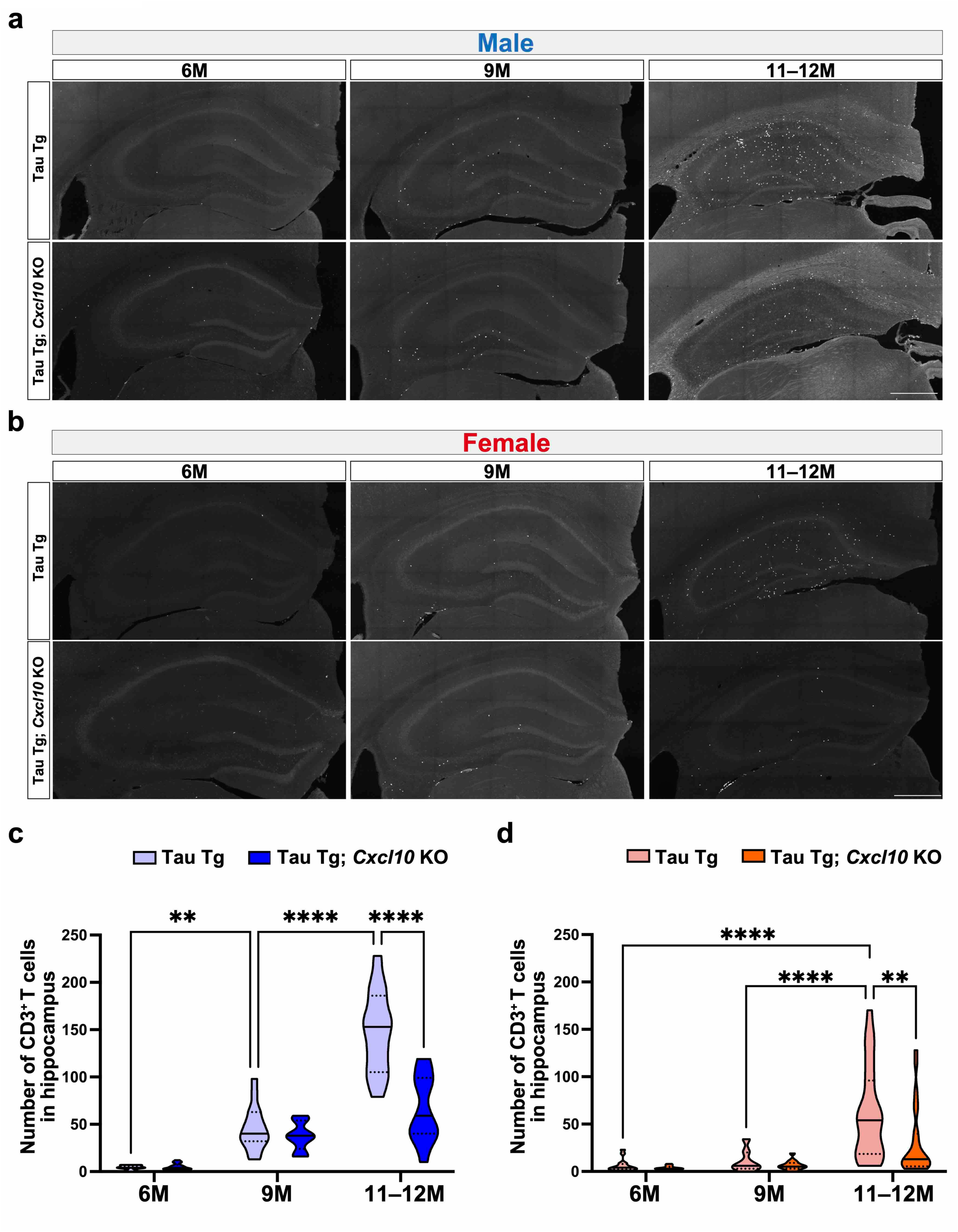
Reduced number of brain T cells in 11–12-month-old Tau Tg mice of both sexes due to *Cxcl10* deficiency. a, **b** Representative images of CD3^+^ T cells in the hippocampi of male (**a**) and female (**b**) Tau Tg and Tau Tg; *Cxcl10* KO mice aged 6 to 11–12 months. Scale bar = 500 μm. **c, d** Quantification of hippocampal T cells in Tau Tg and Tau Tg; *Cxcl10* KO mice aged 6 to 11–12 months (**c**: male, **d**: female). Data are presented as violin plots (n = 15–24 sections from 5–8 mice per group). Statistical analysis was performed using a two-way repeated measures ANOVA followed by a Šídák’s multiple comparisons test (**c**) and a mixed-effects analysis followed by a Šídák’s multiple comparisons test (**d**). \*\**p* < 0.01, \*\*\*\**p* < 0.0001. Source data are provided in the Source Data file.

Finally, we examined whether the cellular response adjacent to *Cxcl10*^+^ versus *Cxcl10*^-^astrocytes differed between sexes by analyzing DEGs in their respective neighboring cells. We found that male Tau Tg mice significantly upregulated the expression of inflammation-associated genes compared to female mice, regardless of the proximity to *Cxcl10*^+^ cells (Fig. S18a, b). This result suggests that male Tau Tg mice exhibit more inflammatory responses due to factors distinct from the influence of CXCL10. In contrast, our findings suggest that CXCL10 acts as a specific modulator of the microenvironment, activating inflammatory responses and mitigating tau pathology exclusively in females. Based on these observations, we speculate that while various inflammatory factors contribute to tau pathology progression, CXCL10 may play a central role in shaping the inflammatory response in females. Nevertheless, the precise molecular and cellular mechanisms underlying these sex-dependent contributions of CXCL10 to tau pathology remain to be fully elucidated and require further research.

## Discussion

In this study, we have identified CXCL10 as a female-specific regulator of tau pathology progression. Although CXCL10 was upregulated in tau-affected brain regions in both sexes, its genetic deletion selectively attenuated pathological tau accumulation and prolonged survival period in females, without altering gliosis, neurodegeneration, or motor dysfunction. These findings indicate that CXCL10 modulates disease progression rather than acts as a ubiquitous driver of neuroinflammation.

Spatial transcriptomic and histological analyses revealed that CXCL10 is predominantly expressed in pathology-associated astrocytes, whereas its receptor CXCR3 is localized to T cells in the brain parenchyma. We observed *Cxcr3*^+^ T cells in close proximity to *Cxcl10*^+^ astrocytes in the hippocampus, and inflammatory gene programs were clearly enriched in CXCL10-high microenvironments. Consistent with its known chemotactic function, *Cxcl10* deficiency reduced the overall number of brain T cells. However, this reduction did not fully ameliorate tau pathology, suggesting that either a threshold of T cell depletion is required or that pathogenic T cell subsets are maintained independently of CXCL10 signaling. Further characterization of these specific subsets will be essential to clarify their distinct contributions to tau progression.

The upstream mechanisms driving CXCL10 upregulation in tauopathy remain to be fully elucidated. As *Cxcl10* is a canonical interferon-stimulated gene, particularly responsive to IFN-γ, we investigated the involvement of interferon signaling. IFN-γ protein levels have been reported to increase in tauopathy models^27)^, and our analysis demonstrated activation of interferon-responsive pathways in *Cxcl10*^+^ astrocytes compared with *Cxcl10*^-^ astrocytes (Fig. S19a, b). These findings support a hypothesis in which IFN-γ signaling is a primary driver of CXCL10 upregulation within pathology-associated astrocytes.

A central finding of this study is the female-specific effect of CXCL10 on tau pathology. Sex hormones are often regarded as primary factors underlying biological sex differences, and have been shown to regulate CXCL10 expression in peripheral tissue culture systems^28)^. However, because CXCL10 expression levels did not differ here between mice of different sexes, the involvement of sex hormones in this context may be minimal. Moreover, differential ligand availability is unlikely to explain this dimorphism. Instead, sex-dependent differences in responsiveness to downstream CXCL10-CXCR3 signaling may underlie the observed phenotype. Notably, *Cxcr3* is encoded on the X chromosome and may escape X chromosome inactivation in certain contexts^29)^, potentially enhancing signaling capacity in females. Alternatively, inflammatory cascades triggered by CXCL10 may exert a proportionally greater influence on disease progression in the female brain. Elucidating the molecular and cellular basis of this sexual dimorphism will be essential for understanding the interface between immune signaling and tauopathy.

Collectively, our results raise several important questions for future investigation. First, this study relied on a single tauopathy model, and the generalizability of CXCL10-dependent mechanisms across distinct tauopathies remains to be established. Furthermore, the precise molecular and cellular pathways linking CXCL10 signaling to tau phosphorylation and aggregation require further clarification.

In summary, our findings identify CXCL10 as a sex-specific immune mediator that shapes the inflammatory microenvironment and promotes tau pathology. Targeting the CXCL10-CXCR3 axis may therefore offer a strategy for addressing sexually dimorphic mechanisms of tau progression. Furthermore, the aspects identified in this study highlight the inherent diversity in the progression patterns of tau pathology. This complexity underscores the necessity of a multifaceted therapeutic approach—either by targeting specific pathways at the optimal temporal window or by simultaneously modulating multiple regulatory points—to effectively halt the progression of tauopathies.

## Methods

### Animals

Animal experiments were approved by the Institutional Animal Care and Use Committee of the Nagoya City University (approval number: 23-015) and conducted in accordance with the Nagoya City University’s guidelines and regulations. PS19 mice were provided by Drs. John Q Trojanowski and Virginia M.-Y. Lee at University of Pennsylvania School of Medicine (RRID:IMSR_JAX:008169)^30)^. These mice express P301S-mutant human microtubule-associated protein tau (MAPT) and exhibit the prominent expression of phosphorylated and insoluble tau protein. In this study, PS19 mice were back-crossed onto a C57BL/6J genetic background, with the resultant line called Tau Tg mice. B6.129S4-*Cxcl10^tm1Adl^*/J (*Cxcl10* KO, RRID:IMSR_JAX:006087) mice and B6.129X1-*Mapt^tm1Hnd^*/J (*Mapt* KO, RRID:IMSR_JAX:007251) mice were obtained from the Jackson Laboratory (Bar Harbor, ME, USA). Animals were housed in rooms maintained at 22–25℃ and 20–60% humidity with an 08:00 (lights on) – 20:00 (lights off) light cycle, and allowed ad libitum access to food and water. All experiments were performed in a blinded manner so as not to let the experimenter know specific details about the mice.

### Sample preparation

Mice were deeply anesthetized by intraperitoneal injection of 0.75 mg/kg medetomidine, 4 mg/kg midazolam, and 5 mg/kg butorphanol. They were then perfused transcardially with phosphate-buffered saline (PBS) and subsequently euthanized in accordance with institutional and national guidelines. After decapitation and removal of the entire brain except for the olfactory bulb and cerebellum, the right and left hemispheres were collected separately. The left hemisphere was immersed overnight at 4°C in freshly prepared 4% formaldehyde made from paraformaldehyde. The tissue was then transferred to 15% sucrose and stored at 4°C until embedding with O.C.T. compound. Coronal brain sections (thickness 30 μm) were prepared using a cryostat. Some tissue samples were embedded in paraffin. The hippocampus and cerebral cortex were isolated from the right hemisphere, snap-frozen in liquid nitrogen, and stored at-80°C

### RNA-seq

Total RNA was extracted from the hippocampi of three 12-month-old male Tau Tg mice and three age-matched WT mice using TRIzol® Reagent (Thermo Fisher Scientific, 15596026, Waltham, MA, USA). The extracted RNA was further purified using the RNeasy Plus Mini Kit (QIAGEN, 74134, Hilden, Germany). Bulk RNA Barcoding and sequencing (BRB-seq)^31)^ was performed for preparing libraries, with some modifications as described in the following. Oligo-dT-based primer was used for single-stranded synthesis, and a Second Strand Synthesis Module (NEB, E6111, Ipswich, MA, USA) was used for double-stranded cDNA synthesis. In-house MEDS-B Tn5 transposase^32)^ was used for tagmentation and amplified by 10 cycles of PCR using Phusion High-Fidelity DNA Polymerase (Thermo Scientific, M0530). Eighty-one base pairs (bp) of insert read (Read2) were obtained on an Illumina NovaSeq 6000 next generation sequencer. For downstream analysis, differentially expressed genes (DEGs) were extracted with DESeq2 (1.42.0) R package software using |log2FC| > 1 and adjusted p-value (padj) < 0.1 as threshold values. We then conducted PCA, heatmap analysis, GSEA, and GO pathway analysis using R. We used MH: Hallmark gene sets in the Mouse Molecular Signatures Database for GSEA analysis.

### Protein array analysis and ELISA

Cytokine and chemokine quantification in brains was performed using brain tissue lysates. Total protein concentrations of lysates were determined by bicinchoninic acid assay, with the samples adjusted to contain equal amounts of protein. Cytokine and chemokine array analyses were performed using the Proteome Profiler Mouse Cytokine Array Kit, Panel A (R&D Systems, ARY006, Minneapolis, MN, USA). CXCL10 protein was quantified using the Mouse IP-10 (CXCL10) ELISA Kit (Thermo Fisher, BMS6018) according to the manufacturer’s instructions.

### Extraction of sarkosyl-insoluble tau

Sarkosyl-insoluble tau was prepared as described by Sahara *et al*^33)^. Briefly, the cerebral cortex, including the piriform and entorhinal cortices, was homogenized in 30 volumes of buffer A (50 mM Tris-HCl, pH 8.0; 274 mM NaCl; 5 mM KCl; and 1 mM phenylmethanesulfonyl fluoride) supplemented with a protease inhibitor cocktail (Roche, 6538304001, Basel, Switzerland) and a phosphatase inhibitor cocktail (Sigma-Aldrich, P5726, St. Louis, MO, USA). The homogenates were centrifuged at 26,300 × g for 20 minutes at 4℃. The resulting pellets were resuspended in 15 volumes of buffer B (10 mM Tris-HCl, pH 7.4; 0.8 M NaCl; 10% sucrose; 1 mM ethylene glycol tetraacetic acid; and 1 mM phenylmethanesulfonyl fluoride) and centrifuged again at 26,300 × g for 20 minutes at 4℃. The resulting supernatants were incubated with 1% sarkosyl for 1 hour at 37℃. Following incubation, the mixture was centrifuged at 150,000 × g for 1 hour at 4℃. The final pellets were resuspended in 1.5 volumes of TE buffer (10 mM Tris-HCl, pH 8.0; 1 mM ethylenediaminetetraacetic acid).

### Western blot

Brain tissues were homogenized in lysis buffer (20 mM Tris-HCl, pH 7.4; 1% Triton X-100) supplemented with a protease inhibitor cocktail and a phosphatase inhibitor cocktail using a Multi-beads shocker. Tissue lysate (15 µg of protein) or 10 µL of sarkosyl-insoluble fraction were electrophoresed on 10% polyacrylamide gels containing 2,2,2-trichloroethanol, which enables visualization of total protein^34)^. After transferring to polyvinylidene difluoride membranes, the membranes were incubated with 2% ECL Prime Blocking Agent (Cytiva, RPN418, Marlborough, MA, USA) and then with primary antibodies at 4℃ overnight. Blots were developed using horseradish peroxidase-conjugated secondary antibodies and ECL reagents (Start, Prime, or Select), and images were captured with a ChemiDoc Touch MP Imaging System. Band intensities were measured and normalized to total protein levels visualized by Stain-Free technology using Image Lab software. Detailed information about the antibodies and equipment used are listed in Supplementary Tables 1 and 4.

### Immunohistochemistry

Brain sections were mounted on glass slides and antigen retrieval was performed using 10 mM citrate buffer (pH6.0) at 121℃ for 5 minutes. The sections were then incubated with blocking buffer (3% donkey serum/PBS containing 0.3% Triton X-100), followed by primary antibodies (anti-CD3, CD31, or NeuN) at 4℃ overnight. Thereafter, the sections were incubated with fluorophore-conjugated secondary antibodies at room temperature for 2 hours, stained with 0.1% Sudan black, and mounted using Fluoromount-G Mounting Medium (Thermo Fisher, 00-4958-02). Free-floating brain sections were stained with anti-Iba1 and anti-C3 antibodies. For Iba1 staining, antigen retrieval was performed with 1% sodium dodecyl sulfate prior to incubation with blocking buffer and primary antibodies at 4°C overnight, whereas C3 staining was performed without antigen retrieval using the same incubation procedure. The sections were then incubated with fluorophore-conjugated secondary antibodies at room temperature for 2 hours and stained with 0.1% Sudan Black, followed by mounting. Immunostaining with the anti-phosphorylated tau [AT8] antibody was visualized using the 3,3′-Diaminobenzidine (DAB)-Nickel method. Briefly, antigen-retrieved, glass-mounted brain sections were incubated with 0.3% hydrogen peroxide for 10 minutes to inactivate endogenous peroxidase, followed by incubation with the anti-phosphorylated tau antibody [AT8] at 4°C overnight. The sections were then incubated with a biotinylated secondary antibody and subsequently with ABC staining solution according to the manufacturer’s protocol (VECTASTAIN Elite ABC Standard Kit, PK-6100, Vector Laboratories, Burlingame, CA, USA), followed by development with DAB-Nickel solution. Finally, the sections were dehydrated, cleared, and coverslipped with Entellan New (Merck Millipore, 501-05301, Darmstadt, Germany). Sections stained with AT8 antibody were imaged using a NanoZoomer S60 virtual slide scanner (NanoZoomer S60, Hamamatsu Photonics, Japan) at ×40 magnification, and the immunopositive area in each image was quantified using FIJI software.

Images of CD3 and CD31 co-staining were acquired using a confocal laser scanning microscope (FV3000, EVIDENT, Tokyo, Japan) with a ×20 objective lens. Tiling was performed to visualize the entire hippocampus. CD3-positive and CD31-negative T cells in the brain parenchyma of the hippocampus were counted using QuPath software. Sections stained with anti-Iba1 and anti-C3 antibodies were imaged using a FV3000 microscope, and the immunopositive area in individual images was quantified using FIJI software. NeuN-stained sections and corresponding bright-field images were captured using the NanoZoomer S60 virtual slide scanner at ×40 magnification. NeuN-immunopositive cells in the CA1, CA3, and DG layers of the hippocampus were measured by using QuPath software. Brain atrophy was evaluated using bright-field images of serial brain sections (spaced 300 µm apart) ranging from bregma-0.95 mm to-3.79 mm with QuPath software according to a previously published method with minor modifications^35)^. Details of antibodies used are listed in Supplementary Table 3 and 4.

### Behavior analysis

Mice were transferred to the test room at least one week prior to the start of behavioral testing to adapt them to the environment and lighting conditions. The temperature of the test room was maintained at 22–25 °C.

**Open field test:** Data collection and analysis were performed using Smart 3.0 video tracking software (PanLab, Barcelona, Spain). Mice were placed in the center of a dimly lit (127–139 lux) open-field arena (45 × 45 × 45 cm) and allowed to freely explore for 10 minutes. The arena was divided virtually into a 4 × 4 grid, with the central 2 × 2 area defined as the center zone. The following parameters were automatically recorded: total distance traveled in the center zone (cm), percentage of total distance in the center zone (%), and mean speed excluding immobility (s). Velocity was calculated by dividing the total distance by the mean speed excluding immobility (cm/s).

**Rotarod test:** The rotarod test was performed based on a previously described protocol with slight modifications^36)^. Briefly, mice were trained to balance on a rotarod rotating at a constant speed of 4 rpm for 1 minute. They were then tested on an accelerating rotarod (from 4 to 40 rpm over 5 minutes). The test was performed three times a day for three consecutive days. The latency to fall was recorded for each trial, and the average of three trials was used for analysis. A rest period of at least 30 minutes was provided between trials in a day.

**Grip strength test:** Forelimb and all-limb grip strength tests were performed using previously described methods^37)^. Grip strength was measured with a Grip Strength Meters for Rats & Mice (Muromachi Kikai, Tokyo, Japan). Mice were held by the tail and allowed to grasp a metal grid with either their forelimbs or all four limbs. They were then gently pulled backward in a horizontal direction, and the peak force was recorded. Each mouse underwent two trials, with the mean value used for analysis. A rest period of at least 5 minutes was provided between trials.

**Foot clasping test:** The test was performed by suspending the mouse by its tail and recording their postures for 30 seconds. The hind limb clasping score was assessed as follows: a score of 0 was assigned if both hind limbs remained extended for more than half of the 30-second period. A score of 1 was given if one hind limb remained extended for more than 15 seconds. A score of 2 was assigned if both hind limbs were in contact with the abdomen for more than 15 seconds. If the hind limbs were clasped and crossed in front of the abdomen, a score of 3 was given.

### Spatial transcriptome analysis by Xenium

Brain hemispheres were prepared as described in the section’Sample preparation,’ with the exception that RNase-free PBS (-) was used for transcardial perfusion. FFPE brain sections were analyzed using Xenium spatial transcriptome analysis (10x Genomics). Sample processing, imaging, and downstream data analysis were performed by KOTAI Biotechnologies. In brief, the filtered data were preprocessed, which included lognormalization with NormalizeData, feature selection with FindVariableFeatures, scaling with ScaleData, PCA with RunPCA (npcs=30), and UMAP with RunUMAP (dims=1:30). The Seurat R toolkit (v5.3.0) was used. For clustering, a nearest-neighbor graph was constructed using FindNeighbors, and a graph-based community detection was performed using FindClusters. The Harmony integrated Xenium data was integrated with publicly available scRNA seq data. The integration was performed using the doublet mode of RCTD deconvolution method in spacexr v2.2.1 with UMI_min = 100 (default). After integration and annotation, we further re-clustered the AC-Epen and immune clusters, respectively (Resolution of re-clustering AC-Epen and Imune = 0.2 and 0.4). For downstream analysis, DEGs were extracted using the Wilcoxon rank sum test. GO analysis data was analyzed using a maximum 500 top up or down regulated genes with padj < 0.05. UMAP and spatial information data figures were produced using Loupe Browser 9.0 and Xenium Explorer4.1.1.

### RNAscope

Fluorescence *in situ* hybridization was performed using the RNAscope Multiplex Assay V2 Kit (Advanced Cell Diagnostics [ACD], 323100, Newark, CA, USA) according to the manufacturer’s instructions with minor modifications. Paraffin-embedded brain tissues were sectioned at 5 µm, mounted on glass slides, dewaxed, treated with hydrogen peroxide, and boiled for 15 minutes in Target Retrieval Reagent (ACD, 322000). After incubation with Protease Plus (ACD, 322331) at 40℃ for 30 minutes, specimens were hybridized with *Cxcl10* probe (408921-C2) at 40°C for 2 hours, followed by signal amplification. Signals were visualized with Alexa Fluor™ 488 Tyramide Reagent (Thermo Fisher, B40953). Slides were counterstained with 4’,6-diamidino-2-phenylindole (DAPI), incubated with a 0.1% Sudan Black, and mounted with Prolong^TM^ Glass Antifade Mountant (Thermo Fisher, P36982).

For co-staining with anti-GFAP, Iba1, and CD3 antibodies, 10 µm-thick cryosections were mounted on glass slides and fixed with 4% formaldehyde. After treatment with H_2_O_2_, the sections were incubated with primary antibodies at 4°C overnight. The sections were then re-fixed with 4% formaldehyde and hybridized with *Cxcl10* probe and *Cxcr3* probe (408921-C2) at 40°C for 2 hours, followed by signal amplification. Signals were visualized with Alexa Fluor™ 488 Tyramide Reagent. Subsequently, the sections were incubated with fluorophore-conjugated secondary antibodies. The remaining procedures were performed as described for the paraffin-embedded samples. Detailed information on the antibodies used is provided in Supplementary Table 2.

### Re-analysis of snRNA-seq and scRNA-seq

SnRNA-seq and scRNA-seq data were re-analyzed using publicly available datasets (GSE218728 and GSE221856, respectively). For snRNA-seq analysis, we used source data from three PS19 mice and three WT mice. Data integration and UMAP visualization were performed using the code provided in the original publication. For downstream analysis, we identified clusters expressing astrocyte-specific marker genes (*Aldh1l1*, *Aqp4*) and microglia-specific marker genes (*Cx3cr1*, *Hexb*) using Feature Plot (Fig. S12b, c). These clusters were extracted using the subset function in the Seurat package (5.4.0). DEGs were identified using the FindMarkers function, and the expression of *Cxcl10* mRNA in each cluster was visualized with the EnhancedVolcano function (version 1.28.2).

For the scRNA-seq analysis, data from PS19 mice with knock-in (KI) of the human APOE4 gene were utilized, with human APOE4 KI mice used as control. Following the referenced study^8)^, we first removed cells with >20% mitochondrial gene content and those with ≥350 detected genes in the scRNA-seq analysis. Each dataset was then normalized using the SCTransform function. Doublets were excluded by the DoubletFinder package (version 2.0.6). For integration, we selected 2,000 features using the SelectIntegrationFeatures function and prepared the data using the PrepSCTIntegration function. Integration anchors were identified with the FindIntegrationAnchors function, and datasets were merged using the IntegrateData function. Dimensionality reduction was performed via PCA, and the first 20 principal components (PCs) were used to generate UMAP with the RunUMAP function. Clustering was conducted using the FindNeighbors and FindClusters functions (resolution = 0.1), based on the same PCs. Cell types were annotated based on canonical marker genes identified using the FindAllMarkers function. Clusters lacking defining markers were excluded from further analysis (Fig. S13a). The expression pattern of *Cxcr3* across identified cell types was visualized using the Feature Plot and Violin Plot functions.

### Flow cytometry

Mice were deeply anesthetized and then perfused with PBS to remove circulating blood. Brain tissues were dissected, and the hippocampus and cerebral cortex were mechanically minced and enzymatically dissociated in Hanks’ Balanced Salt Solution (HBSS) containing 1 mg/mL collagenase type IV (Sigma, C5138) and 0.2 mg/mL DNase I (Roche, 104159) at 37°C for 15 minutes. The tissues were subsequently passed 10 times through 18-gauge and then 21-gauge needles attached to syringes. Mononuclear cells were isolated by density gradient centrifugation at 1,000 × g for 15 minutes using 37%/70% Percoll (GE Healthcare Life Sciences, Uppsala, Sweden). Isolated cells were treated with 5 µg/mL Fc block solution (BD Biosciences, 553141, San Jose, CA, USA) at 4℃ for 10 minutes, followed by staining with fluorophore-conjugated antibodies. Cells were analyzed using a FACS Aria III (BD) and FlowJo software (BD). Detailed information on the antibodies and equipment used is listed in Supplementary Table 3, 4.

## Statistical analysis

All statistical analyses were performed using GraphPad Prism software with a significance threshold of *p* < 0.05. Data were presented as mean ± standard error of mean (S.E.M.). The Shapiro-Wilk test was used to assess the normality of data distribution. The F test or Brown-Forsythe test was applied to evaluate the homogeneity of variances. For comparisons between two groups, an unpaired two-tailed t-test was used when data were normally distributed, but otherwise the Mann-Whitney U test was employed. If normality was confirmed but equal variances could not be assumed, an unpaired t-test with Welch’s correction was performed. For comparisons among three or more groups, one-way analysis of variance (ANOVA) followed by a Tukey’s multiple comparisons test was used when data were normally distributed with equal variances. When the data were normally distributed but exhibited unequal variances, Brown-Forsythe ANOVA tests were applied, followed by Dunnett’s T3 multiple comparison test. Conversely, when the data did not follow a Gaussian distribution, a Kruskal-Wallis test was used, followed by Dunn’s multiple comparisons test. To simultaneously analyze the effects of two independent variables on a dependent variable, mixed-effects analysis followed by either Tukey’s or Šídák’s multiple comparisons test was performed. In addition, the log-rank test was used for Kaplan–Meier survival analysis.

## Data availability

All relevant data are available from the corresponding authors upon reasonable request.

## Acknowledgements

We thank Drs. Virgnia M.-Y. Lee and John Q Trojanowski for providing us with the PS19 mice. We also thank the Transcriptomics Research at the Institute of Bioregulatory Medicine, Kyushu University, within the Department of Transcriptomics, Okawa Laboratory, for conducting the bulk RNA-seq analysis. Technical assistance provided by Ms. Keiko Mutsuura is gratefully acknowledged. We also acknowledge assistance received from the Center for Experimental Animal Science and the Center for Research Equipment at Nagoya City University for enabling us to use research equipment shared as part of the MEXT Project for promoting public utilization of advanced research infrastructure (Program for supporting construction of core facilities; Grant Number JPMXS0441500024). We also thank the Center for Animal Resources and Development at Kumamoto University for technical assistance. Figure 1a, 3a, 4a were created using BioRender (https://app.biorender.com/). This research was supported by AMED (Grant Numbers JP20gm1210010 (T.S.), the JST Moonshot R&D Program (Grant Number JPMJMS2024 (T.S)), JST SPRING Japan (Grant Number JPMJSP2130 (R.U.)), and JP24wm0625303 (T.S.)), JSPS KAKENHI (Grant Numbers JP20H03564 (T.S.), JP24K02354 (T.S.), JP22K06865 (M.H.), JP24K18381 (T.M.), and JP24KJ1888 (R.U.)), the Hori Sciences & Arts Foundation (T.S.), a LEGEND Research Grant 2024 from BioLegend (M.H.), a Grant-in-Aid for Research from Nagoya City University (Grant Number 2021101), a Grant-in-Aid for the Outstanding Research Group Support Program at Nagoya City University (Grant Number 2401101), and a Nanken-Kyoten grant (Grant Number 2024-kokunai 24, Grant Number 2025-kokunai 29) form the Institute of Science Tokyo.

