## Supplementary info for "CXCL10 drives female-specific tau pathology progression and defines sex-dependent vulnerability in tauopathy model mice"

**Contents:**

Supplementary Figures 1–19
Supplementary Table 1–4

#### Supplementary Figures

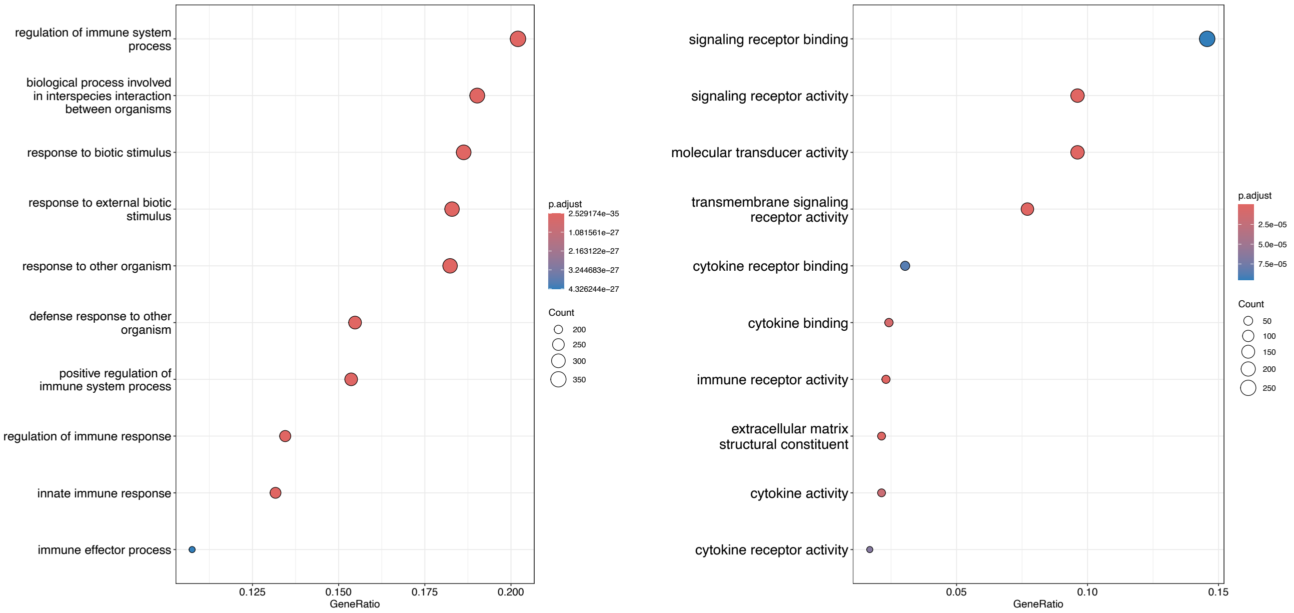


**Supplementary Fig. 1 | Inflammatory response pathway was activated in the hippocampus of Tau Tg mice.**

**a, b** Gene Ontology (GO) pathway analysis focusing on biological processes (BP) and molecular functions (MF) was performed to compare P301S tau transgenic (Tau Tg) mice with wild-type (WT) mice. The top 10 significantly upregulated pathways in Tau Tg mice are shown in each panel. Number of mice used: WT (n = 3), Tau Tg (n = 3). Differentially expressed genes (DEGs) were identified using the Ward test, and GO analysis was evaluated via over-representation analysis employing the hypergeometric test. Source data are provided in the Source Data file.

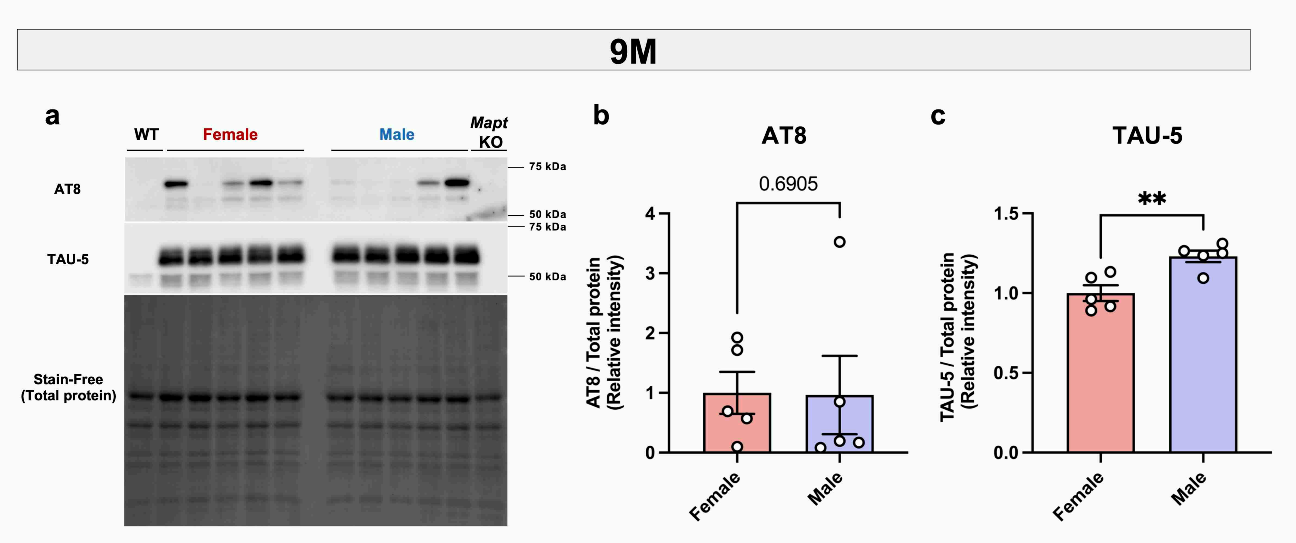


**Supplementary Fig. 2 Sex-dependent changes relative to AT8-positive tau accumulation in 9-month-old Tau Tg mice.**

**a–c** Immunoblot analysis of tau protein in the cerebral cortices of 9-month-old Tau Tg mice of both sexes. Immunoblot images of phosphorylated tau (p-tau) detected with anti-pSer202/pThr205 [AT8] antibodies and total tau detected with TAU-5 antibody (**a**). **b, c** Protein expression levels were normalized to total protein using the Stain-Free method. Data are presented as mean ± S.E.M. Five female and five male Tau Tg mice were used Number of mice used: Female Tau Tg (n = 5), and Male Tau Tg (n = 5). Statistical analysis was performed using an unpaired two-tailed t-test (**c**) or a Mann-Whitney U test (**b**). **$p$ < 0.01. Source data, including full blot images are provided in the Source Data file.

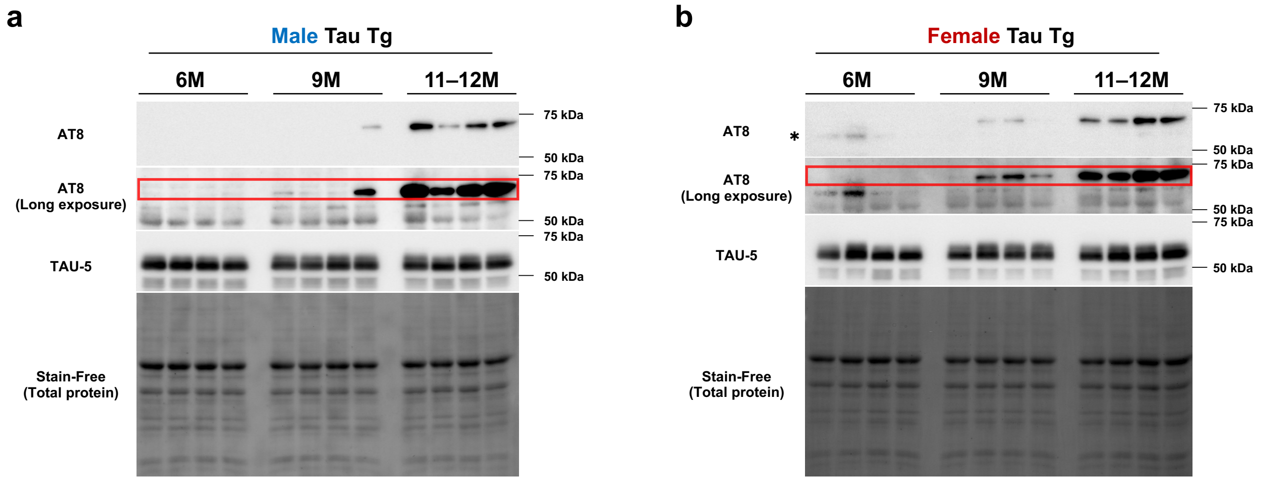


**Supplementary Fig. 3 | Age-dependent changes in AT8-positive tau accumulation.**

**a, b** Immunoblot analysis of tau protein in the cerebral cortices of Tau Tg mice aged 6 to 11–12 months from both sexes. Immunoblot images of p-tau detected with anti-pSer202/pThr205 [AT8] antibodies and total tau detected with TAU-5 antibody (**a**: male, **b**: female). Red rectangle indicates AT8-positive tau band; asterisk denotes a non-specific band. Source data, including full blot images, are provided in the Source Data file.

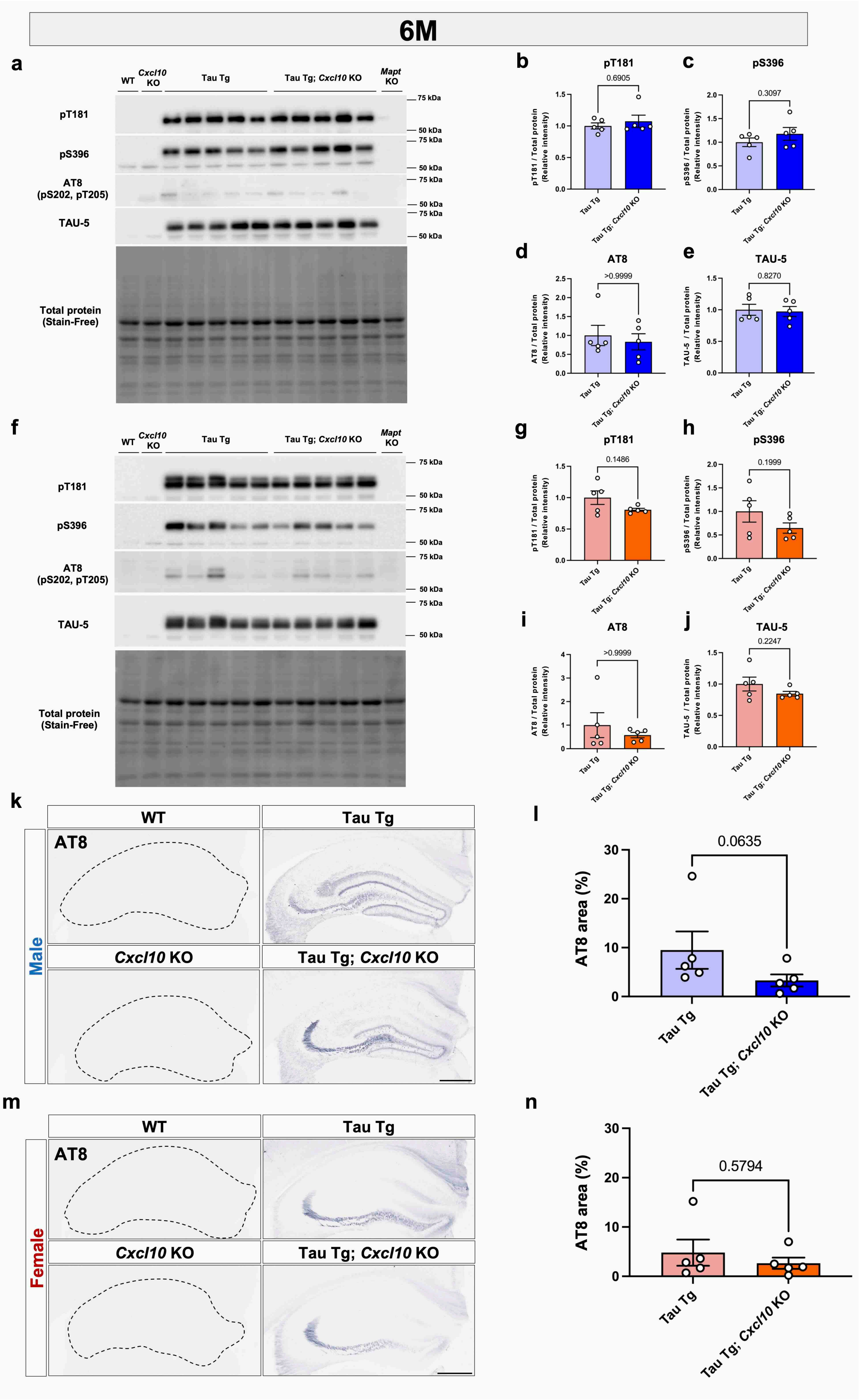
6M
a
WT
Cxcl10 KO
Tau Tg
Tau Tg; Cxcl10 KO
Mapt KO
pT181
pS396
AT8 (pS202, pT205)
TAU-5
Total protein (Stain-Free)
75 kDa
50 kDa
b
pT181
0.6905
c
pS396
0.3097
d
AT8
>0.9999
e
TAU-5
0.8270
f
g
pT181
0.1486
h
pS396
0.1999
i
AT8
>0.9999
j
TAU-5
0.2247
k
WT
Tau Tg
AT8
Male
Cxcl10 KO
Tau Tg; Cxcl10 KO
l
AT8 area (%)
0.0635
m
WT
Tau Tg
AT8
Female
Cxcl10 KO
Tau Tg; Cxcl10 KO
n
AT8 area (%)
0.5794

**Supplementary Fig. 4 | Effect of *Cxcl10* deficiency on tauopathy model mice at 6 months of age**

Immunoblot analysis of tau protein in the cerebral cortices of 6-month-old Tau Tg mice with or without C-X-C motif chemokine ligand 10 (*Cxcl10)* deficiency. Males (**a–e**), females (**f–j**). **a**, **f** Immunoblot images of p-tau detected with anti-pThr181, pSer396, and pSer202/pThr205 [AT8] antibodies, and total tau detected with TAU-5 antibody. **b–e, g–j** Protein expression levels were normalized to total protein using the Stain-Free method. **k, m** Immunohistochemical analysis of p-tau in the hippocampus using AT8 antibody. **l, n** Quantification of the AT8-positive area in males (**l**) and females (**n**). Scale bar = 500 µm. Data are presented as mean ± S.E.M. Number of mice used: Tau Tg (n = 5), and Tau Tg; *Cxcl10* KO (n = 5) for both sexes. Statistical analysis was performed using an unpaired two-tailed t-test (**c**, **e**, **h**, **j**), an unpaired t-test with Welch's correction (**g**), and a Mann-Whitney U test (**b, d, i, l, n**). Source data, including full blot images are provided in the Source Data file.

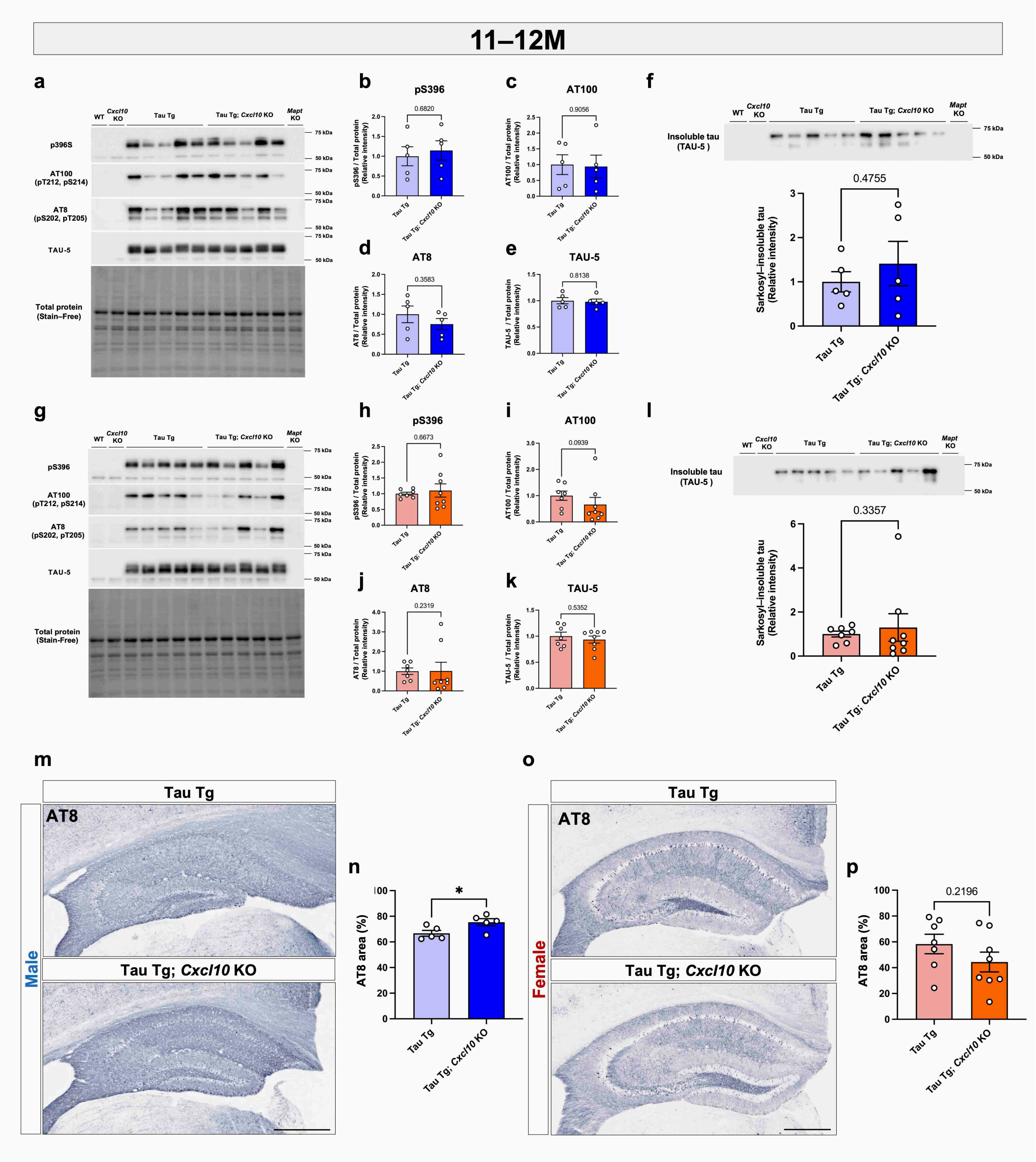
11–12M
a
b
pS396
c
AT100
d
AT8
e
TAU-5
f
Insoluble tau (TAU-5 )
0.4755
Sarkosyl-insoluble tau (Relative intensity)
Tau Tg
Tau Tg; Cxcl10 KO
g
h
pS396
i
AT100
j
AT8
k
TAU-5
l
Insoluble tau (TAU-5 )
0.3357
m
Tau Tg
AT8
Tau Tg; Cxcl10 KO
Male
n
AT8 area (%)
*
o
Female
p
0.2196

**Supplementary Fig. 5 | Biochemical and histological assessment of tau pathology in 11–12-month-old Tau Tg mice with or without *Cxcl10* deficiency.**

Immunoblot analysis of tau protein in the cerebral cortices of 11–12-month-old Tau Tg mice with or without *Cxcl10* deficiency. Males (**a–f, m**), females (**g–l, o**). **a**, **g** Immunoblot images of p-tau detected with anti-pSer396, pThr212/pSer214 [AT100], and pSer202/pThr205 [AT8] antibodies, and total tau detected with TAU-5 antibody. **b**–**e**, **h**–**k** Protein expression levels were normalized to total protein using the Stain-Free method. **f**, **l** Immunoblot images of sarkosyl-insoluble tau from the cerebral cortex, including the piriform and entorhinal cortices. Bar graphs represent relative expression level of tau protein in Tau Tg; *Cxcl10* KO mice compared to Tau Tg mice. **m, o** Immunohistochemical analysis of p-tau in the hippocampus using AT8 antibody. Bar graphs indicate the AT8-positive area in males (**n**) and females (**p**). Scale bar = 500 μm. Data are presented as mean ± S.E.M. Number of mice used: male Tau Tg (n = 5), male Tau Tg; *Cxcl10* KO (n = 5), female Tau Tg (n = 7), and female Tau Tg; *Cxcl10* KO (n = 8). Statistical analysis was performed using an unpaired two-tailed t-test (**b–f, h, k, n, p**) or a Mann-Whitney U test (**i, j, l**). *$p < 0.05$. Source data, including full blot images are provided in the Source Data file**.**

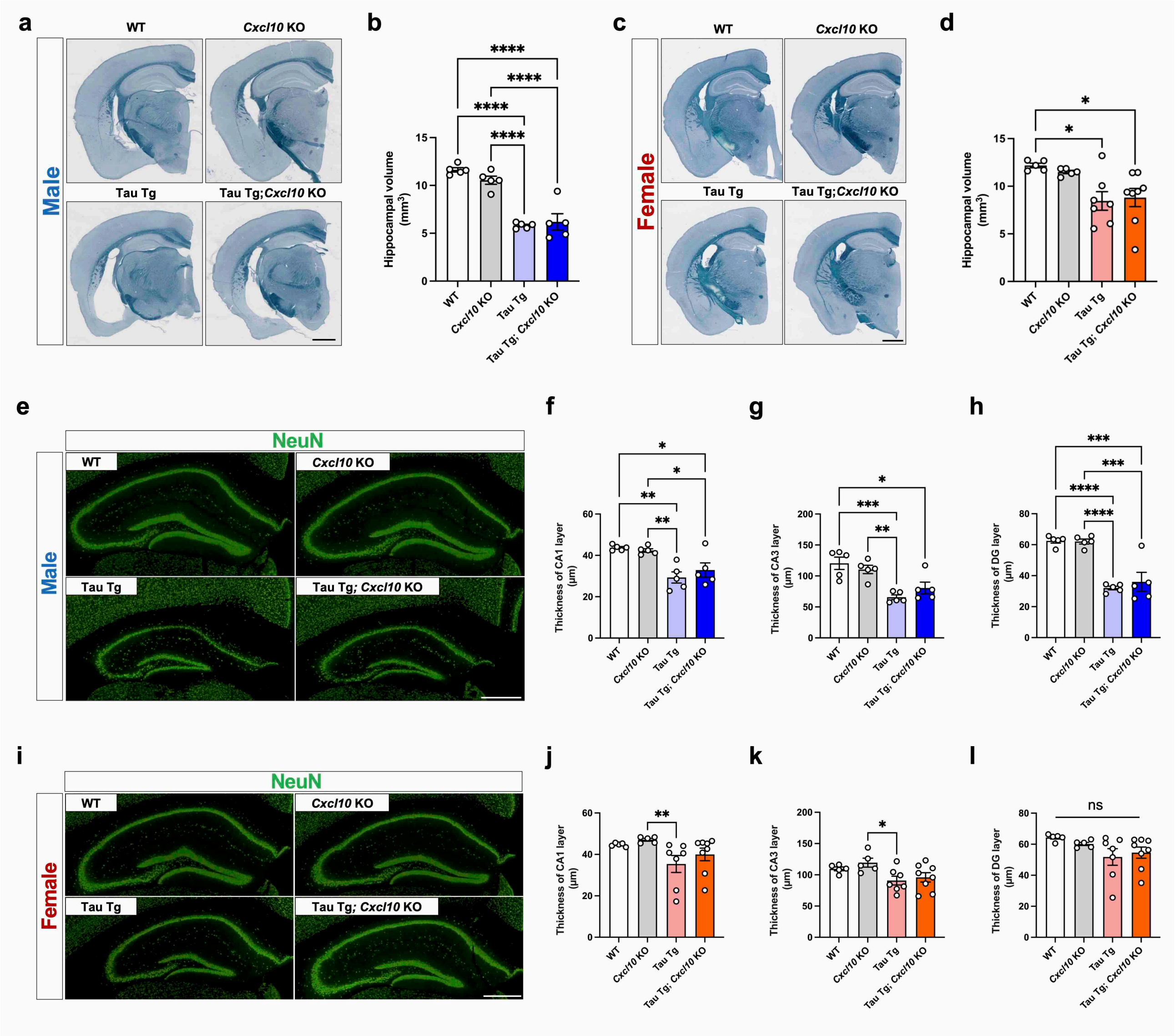


**Supplementary Fig. 6 | Effect of *Cxcl10* deficiency on the brain atrophy, neurodegeneration, and behavior.**

**a, c** Representative images of brain atrophy in age-matched male (**a**) and female (**c**) mice of each genotype are shown. Scale bar = 1 mm. Bar graphs indicate the hippocampal volume in males (**b**) and females (**d**). Data are presented as mean ± S.E.M. **e, i** Representative images of NeuN-stained hippocampal sections from age-matched male (**e**) and female (**i**) mice of each genotype are shown. Scale bar = 500 µm. Bar graphs show the thickness of the CA1 (male: **f**, female: **j**), CA3 (male: **g**, female: **k**), and dentate gyrus (DG) (male: **h**, female: **i**) layers in 11–12-month-old mice. Data are presented as mean ± S.E.M. Number of mice used: male WT (n = 5), male *Cxcl10* KO (n = 5), male Tau Tg (n = 5), and male Tau Tg; *Cxcl10* KO (n = 5). Female WT (n = 5), female *Cxcl10* KO (n = 5), female Tau Tg (n = 7), and female Tau Tg; *Cxcl10* KO (n = 8). Statistical analysis was performed using a one-way ANOVA followed by a Tukey's multiple comparisons test (**b, d, f, g, h, k and l**) and a Kruskal-Wallis test followed by the Dunn's multiple comparisons test (**j**). $^{*}p < 0.05$, $^{**}p < 0.01$, $^{***}p < 0.001$, $^{****}p < 0.0001$. Source data are provided in the Source Data file.

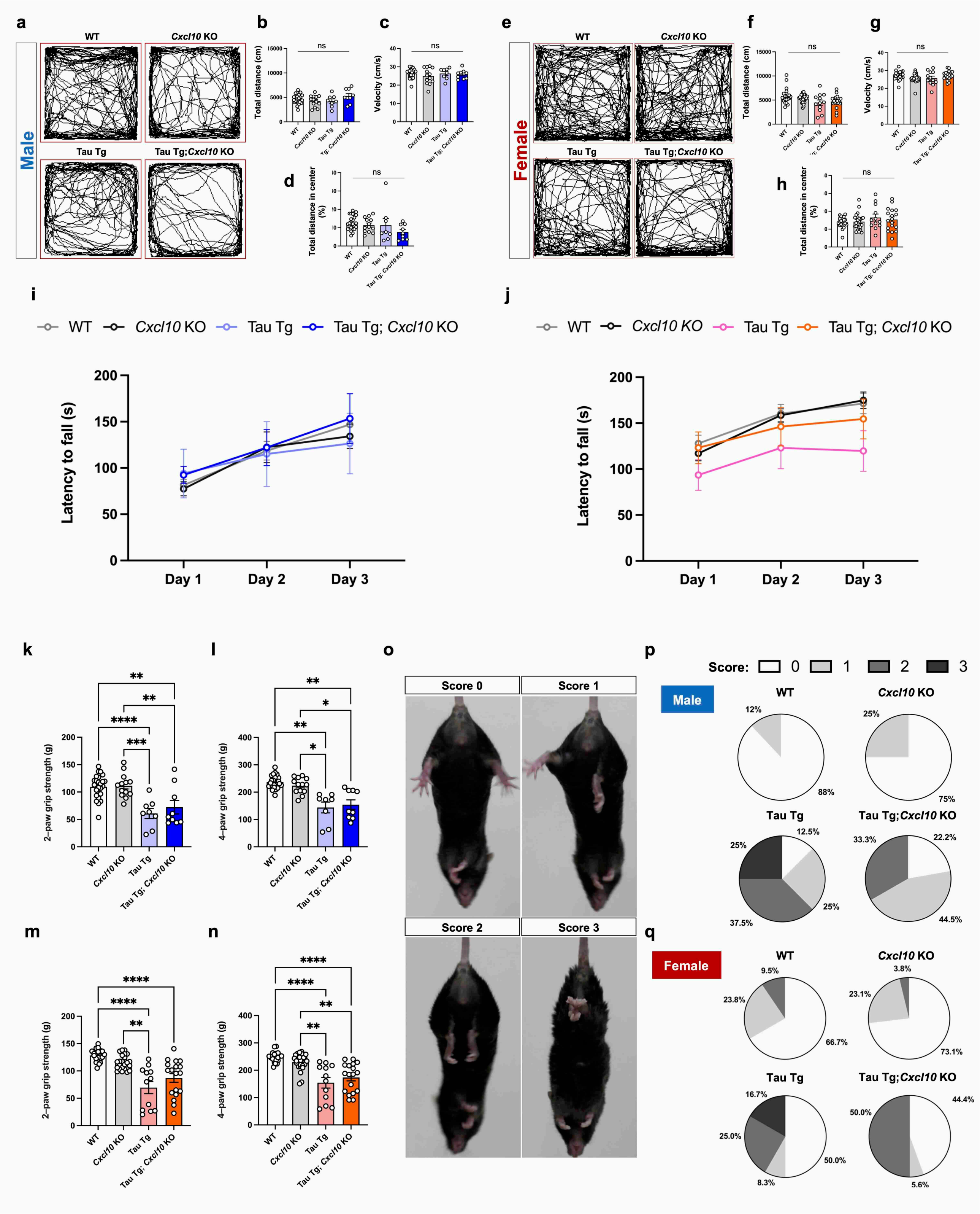

a
WT
Cxcl10 KO
Male
Tau Tg
Tau Tg;Cxcl10 KO
b
c
d
ns
e
Female
f
g
h
i
WT
Cxcl10 KO
Tau Tg
Tau Tg; Cxcl10 KO
Latency to fall (s)
Day 1
Day 2
Day 3
j
k
2-paw grip strength (g)
l
4-paw grip strength (g)
m
n
o
Score 0
Score 1
Score 2
Score 3
p
Score:
Male
Female
q

**Supplementary Fig. 7 | Behavioral assessment of tau pathology-associated 11–12-month-old Tau Tg mice with or without *Cxcl10* deficiency.**

Representative trajectories from the open field test are shown for age-matched male (**a**) and female (**e**) mice of each genotype. **b-d, f-h** Bar graphs show total distance and velocity to assess motor function, while the total distance in the center area is used to evaluate anxiety-like behavior. **i, j** Graphs showing latency to fall for each genotype in mice of both sexes (**i**: male, **j**: female) as measured by the rotarod test. Data are presented as mean ± S.E.M. **k-n** Grip strength tests were conducted for male (**k, l**) and female (**m, n**) mice aged 11–12 months. Results for the two-paw test are shown in (**k, m)**, and the four-paw test are shown in (**l, n**). Data are presented as mean ± S.E.M. Number of mice used: male WT (n = 25), male *Cxcl10* KO (n = 13), male Tau Tg (n = 8), and male Tau Tg; *Cxcl10* KO (n = 9). Female WT (n = 21), female *Cxcl10* KO (n = 26), female Tau Tg (n = 12), and female Tau Tg; *Cxcl10* KO (n = 18). Statistical analyses were performed using a one-way ANOVA followed by a Tukey's multiple comparisons test (**b–d, k**), a Brown-Forsythe ANOVA test followed a Dunnett's T3 multiple comparisons test (**i**), a Kruskal-Wallis test followed by a Dunn's multiple comparisons test (**f–h, m, n**), a mixed-effects analysis followed by a Šídák's multiple comparisons test (**i**), and a mixed-effects analysis followed by a Tukey's multiple comparisons test (**j**). **o** Representative images highlighting abnormalities observed in the hind limbs of age-matched mice. The pie charts show the proportion of each score for each genotype. Male (**p**); female (**q**). Source data are provided in the Source Data file.

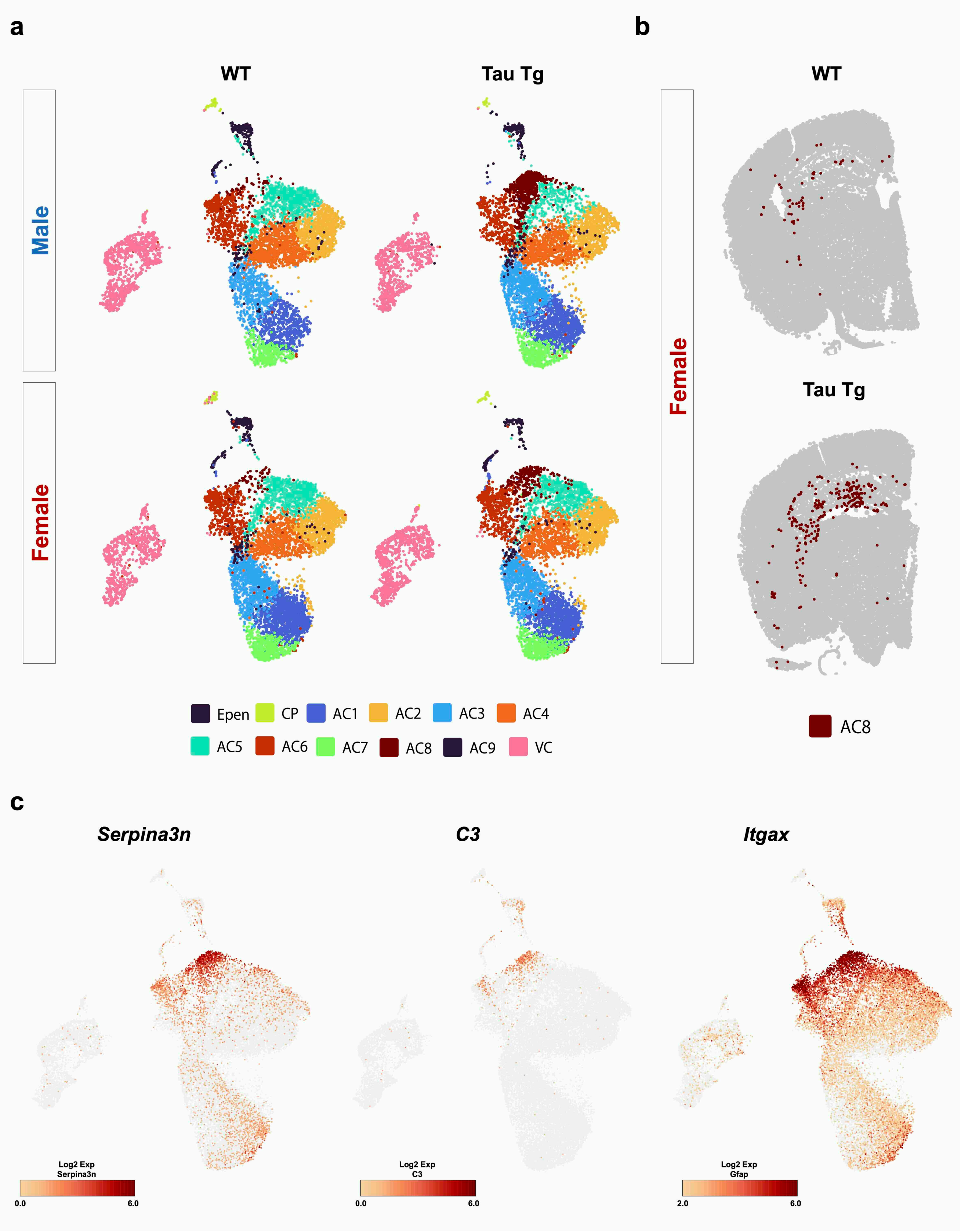
a
b
WT
Tau Tg
Male
Female
WT
Female
Tau Tg
Epen
CP
AC1
AC2
AC3
AC4
AC5
AC6
AC7
AC8
AC9
VC
AC8
c
Serpina3n
C3
Itgax
Log2 Exp
Serpina3n
0.0
6.0
Log2 Exp
C3
0.0
6.0
Log2 Exp
Gfap
2.0
6.0

**Supplementary Fig. 8 | Characterization of AC8 clusters in the brains of Tau Tg mice**

**a** Images classified by genotype and sex for re-clustering of astrocyte-ependymal cells (AC-Epen) cluster from Xenium data. The AC8 cluster was significantly present in the brains of Tau Tg mice regardless of the sex of the animals. **b** Cell localization of AC8 in WT and Tau Tg mouse brains. **c** Projection of selected marker genes for disease-associated astrocyte (*Serpina3n*, *C3*, *Itgax*) in uniform manifold approximation projection (UMAP) by Xenium data. Number of mice used: male WT (n = 1), male Tau Tg (n = 1), female WT (n = 1), and female Tau Tg (n = 1).

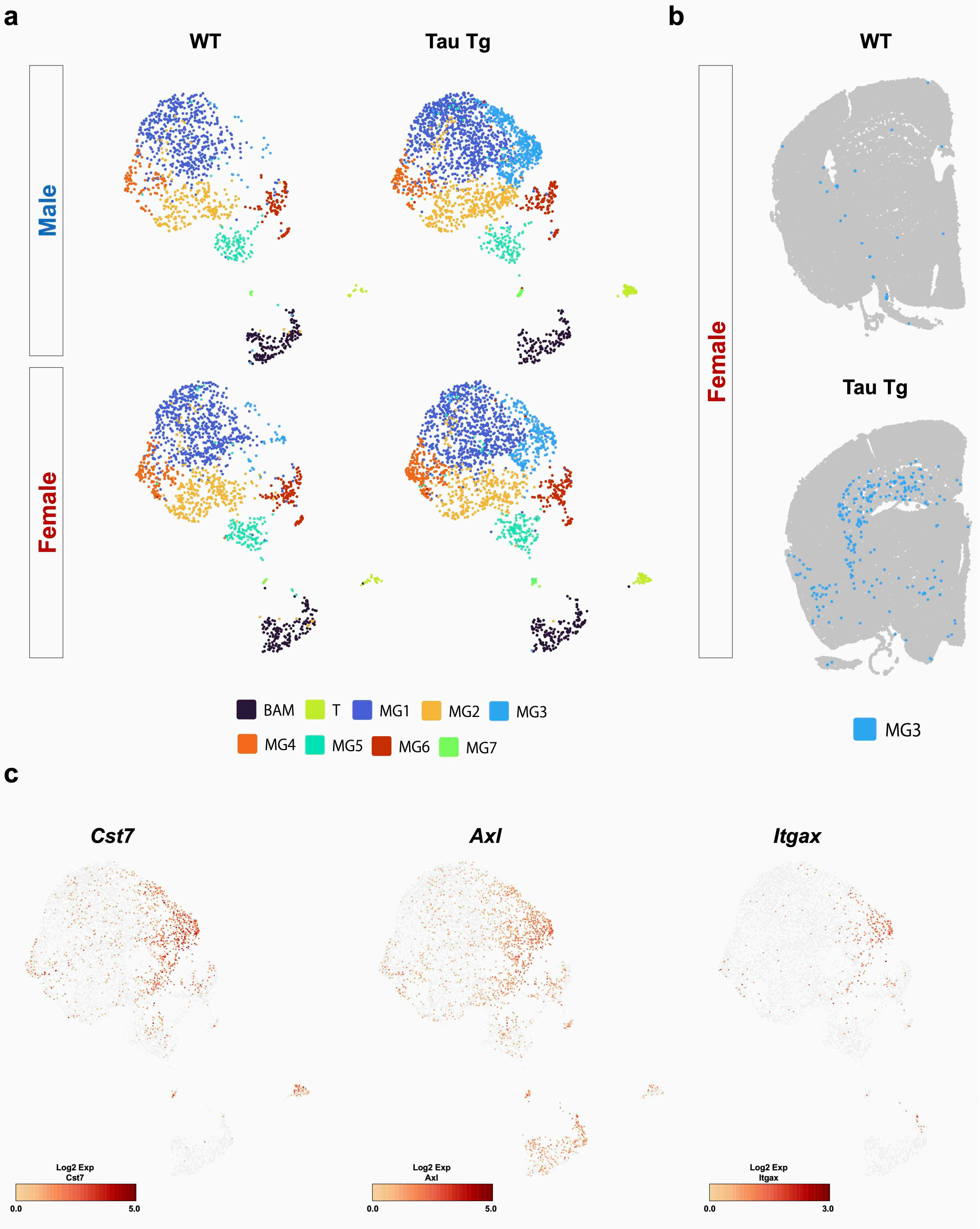

a
WT
Tau Tg
Male
Female
b
WT
Tau Tg
Female
BAM
T
MG1
MG2
MG3
MG4
MG5
MG6
MG7
MG3
c
Cst7
Axl
Itgax
Log2 Exp
Cst7
0.0
5.0
Log2 Exp
Axl
0.0
5.0
Log2 Exp
Itgax
0.0
3.0

**Supplementary Fig. 9 | Characterization of MG3 clusters in the brains of Tau Tg mice**

**a** Images classified by genotype and sex for re-clustering of immune cell clusters from Xenium data. The Microglia 3 (MG3) cluster was significantly present in the brain of Tau Tg mice regardless of the sex of the animals. **b** Cell localization of MG3 in WT and Tau Tg mice brain. **c** Projection of selected marker genes for disease-associated microglia (*Cst7*, *Axl*, *Itgax*) in UMAP from Xenium data. Number of mice used: male WT (n = 1), male Tau Tg (n = 1), female WT (n = 1), and female Tau Tg (n = 1).

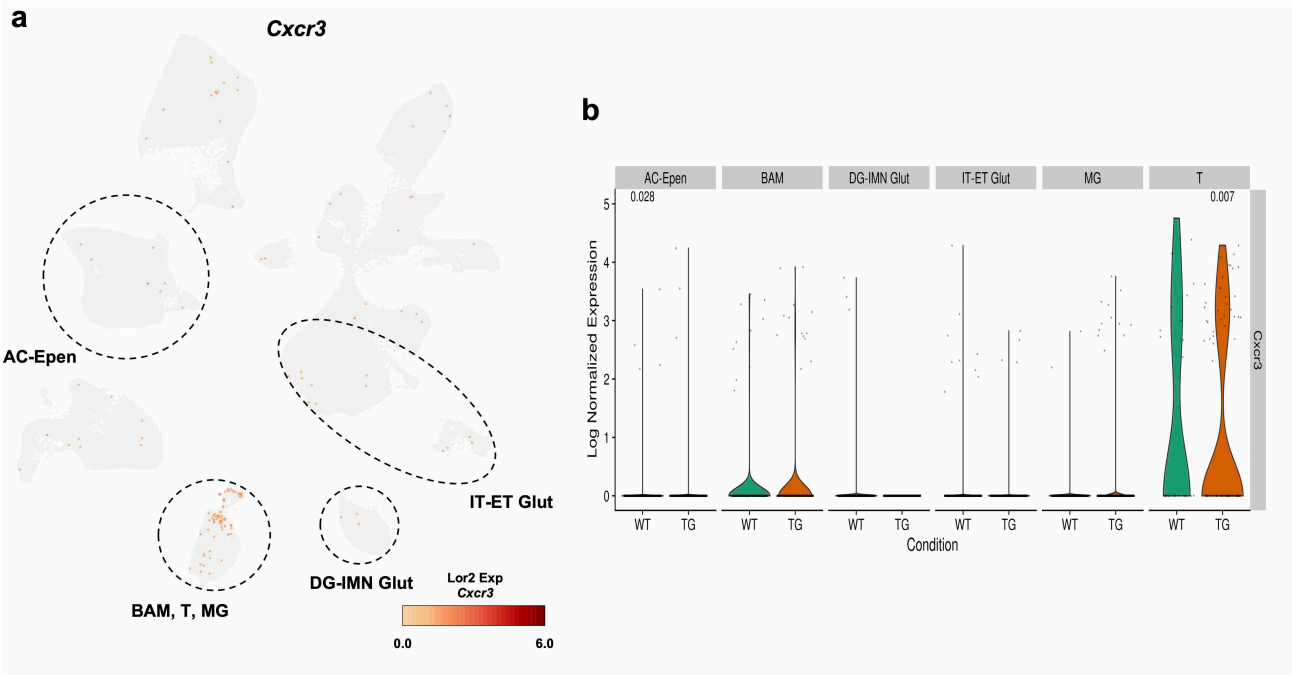


**Supplementary Fig. 10 | Identification of *Cxcr3*-expressing cells in the brains of WT and Tau Tg mice.**

**a** Representative C-X-C motif chemokine receptor 3 (*Cxcr3)* expressing cells in WT and Tau Tg mice. **b** Quantitative *Cxcr3* gene expression using violin plots for AC-Epen, border-associated macrophages (BAM), DG-immature neurons (IMN) glutamatergic (Glut), intratelencephalic (IT)-extratelencephalic (ET) Glut, MG, and T cell (T) types. Statistical analysis was performed using the Wilcoxon test. Number of mice used: WT: male WT (n = 1), female WT (n = 1), TG: male Tau Tg (n = 1), female Tau Tg (n = 1).

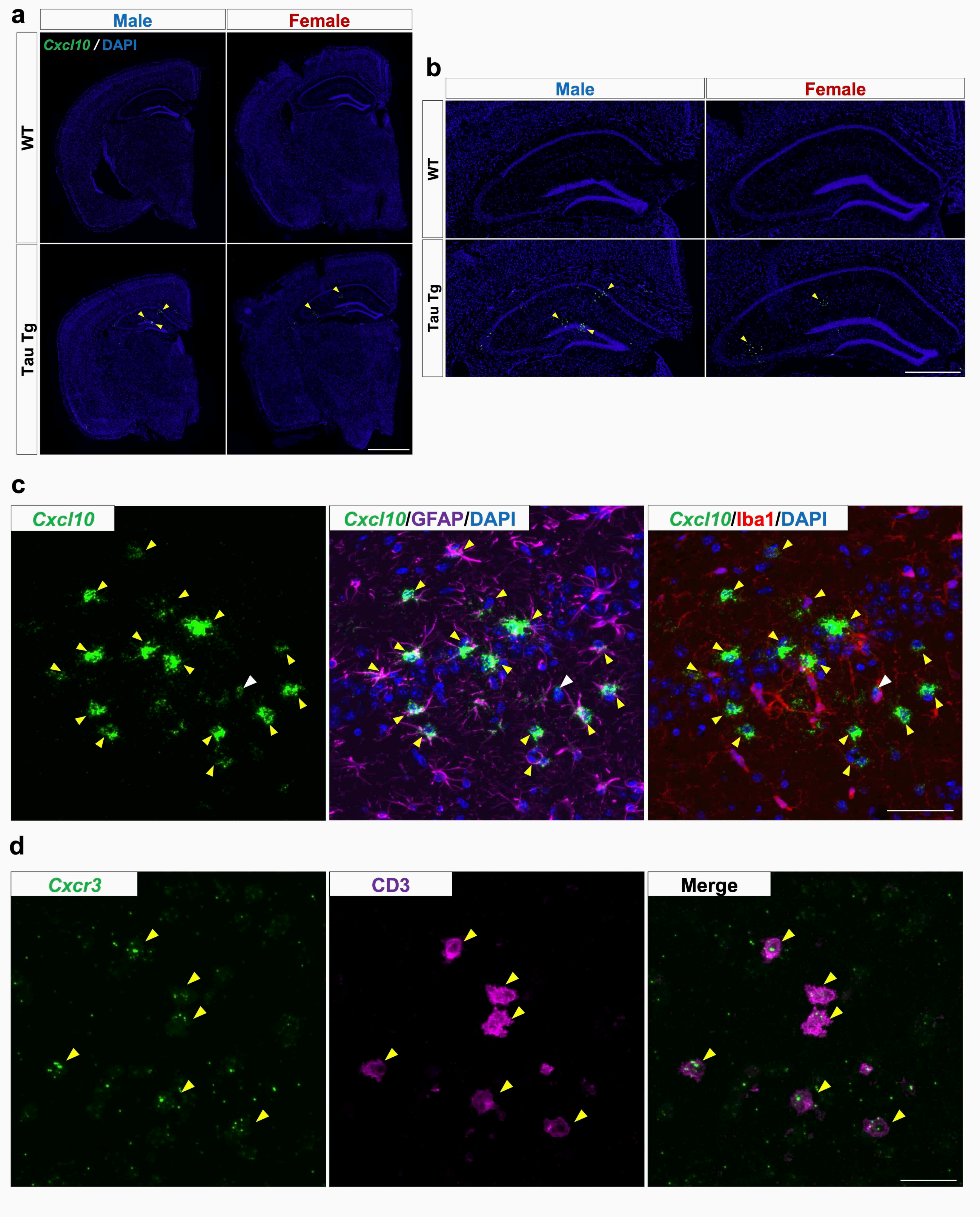
a
Male
Female
Cxcl10 / DAPI
WT
Tau Tg
b
Male
Female
WT
Tau Tg
c
Cxcl10
Cxcl10/GFAP/DAPI
Cxcl10/Iba1/DAPI
d
Cxcr3
CD3
Merge

**Supplementary Fig. 11 | Validation of *Cxcl10* and *Cxcr3*-expressing cells.**

**a** Image showing brain sections stained with RNAscope and DAPI from age-matched WT and Tau Tg mice of both sexes. Yellow arrowheads indicate *Cxcl10*$^{+}$ cells. Scale bar = 1 mm. **b** Images showing enlarged views of the hippocampal sections from **a**. Scale bar = 500 μm. Number of mice used: male WT (n = 1), male Tau Tg (n = 1), female WT (n = 1), and female Tau Tg (n = 1). (**c**, **d**) Images showing *Cxcl10* mRNA- and *Cxcr3* mRNA-positive cells in the hippocampus. *Cxcl10* and *Cxcr3* mRNA were detected using RNAscope probes, while GFAP (an astrocyte marker), Iba1 (a microglial marker), and CD3 (a T cell marker) were identified using Anti-GFAP, Anti-Iba1, and Anti-CD3 antibodies, respectively. The yellow and white arrowheads highlight double-immunopositive cells for *Cxcl10* mRNA and GFAP or Iba1, respectively in (**c**), and *Cxcr3* mRNA and CD3 in (**d**). Scale bar = 50 (**c**) and 20 μm (**d**). Number of mice used: male Tau Tg (n = 1).

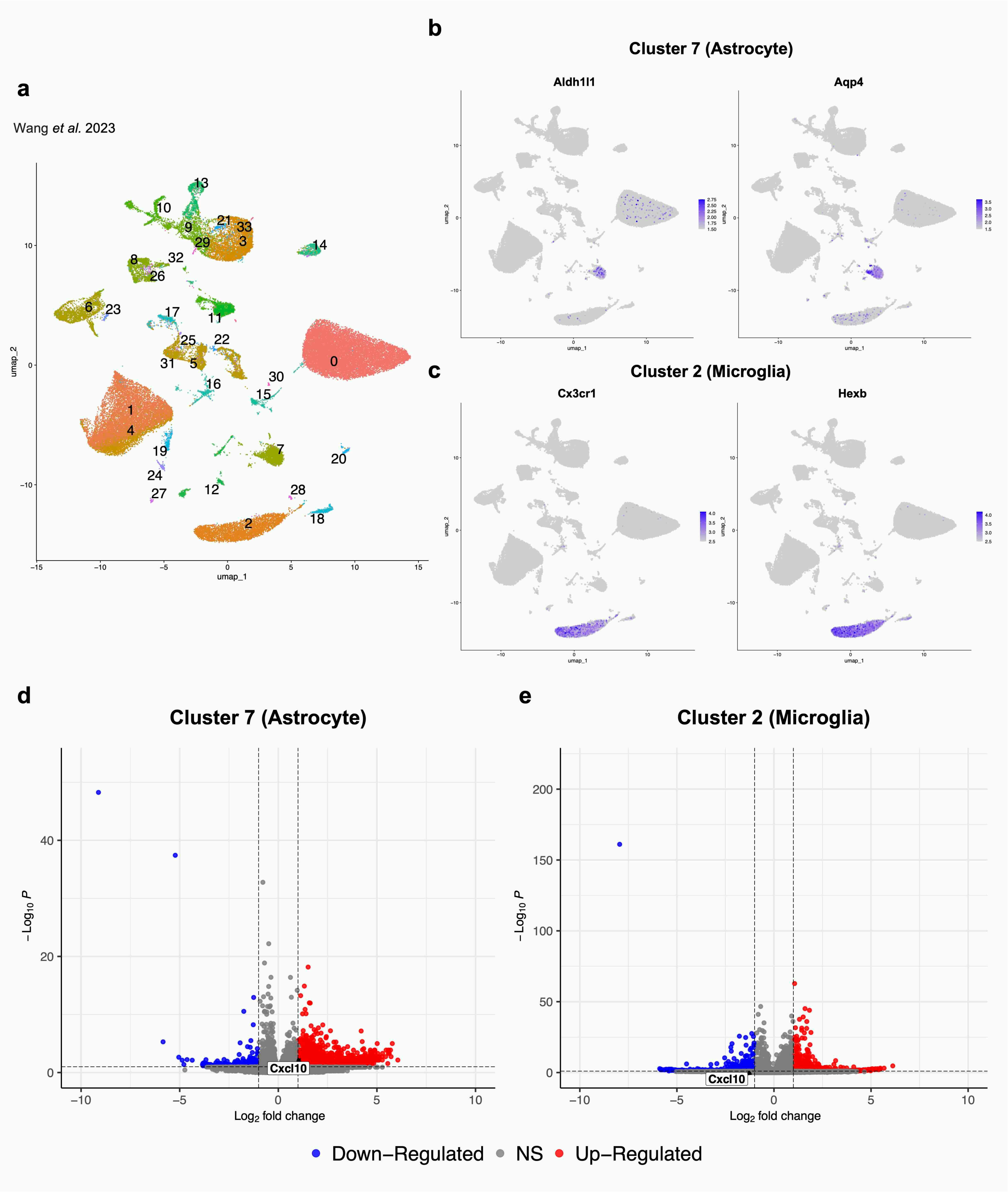


**Supplementary Fig. 12 | Re-analysis of snRNA-seq datasets to identify *Cxcl10*-expressing cell types in tauopathy mouse model.** **a–e** Re-analysis of the single-nucleus RNA sequencing (snRNA-seq) dataset (GSE218728), including samples from three PS19 and three WT mice. **a** UMAP plot showing individual clusters of single nuclei from hippocampus. **b, c** Astrocytic (**b**) and microglial (**c**) clusters were identified using canonical markers. **d, e** $Log_2$ fold change > |1| and p < 0.1 highlighted in red, $Log_2$ fold change < |1| and p > 0.1 shown in blue. All other transcripts are indicated in gray. *Cxcl10* expression in astrocytic (**d**) and microglial (**e**) clusters is displayed in volcano plot. Number of mice used: male WT (n = 2), male PS19 (n = 2), female WT (n = 1), and female PS19 (n = 1). Source data are provided in the Source Data file.

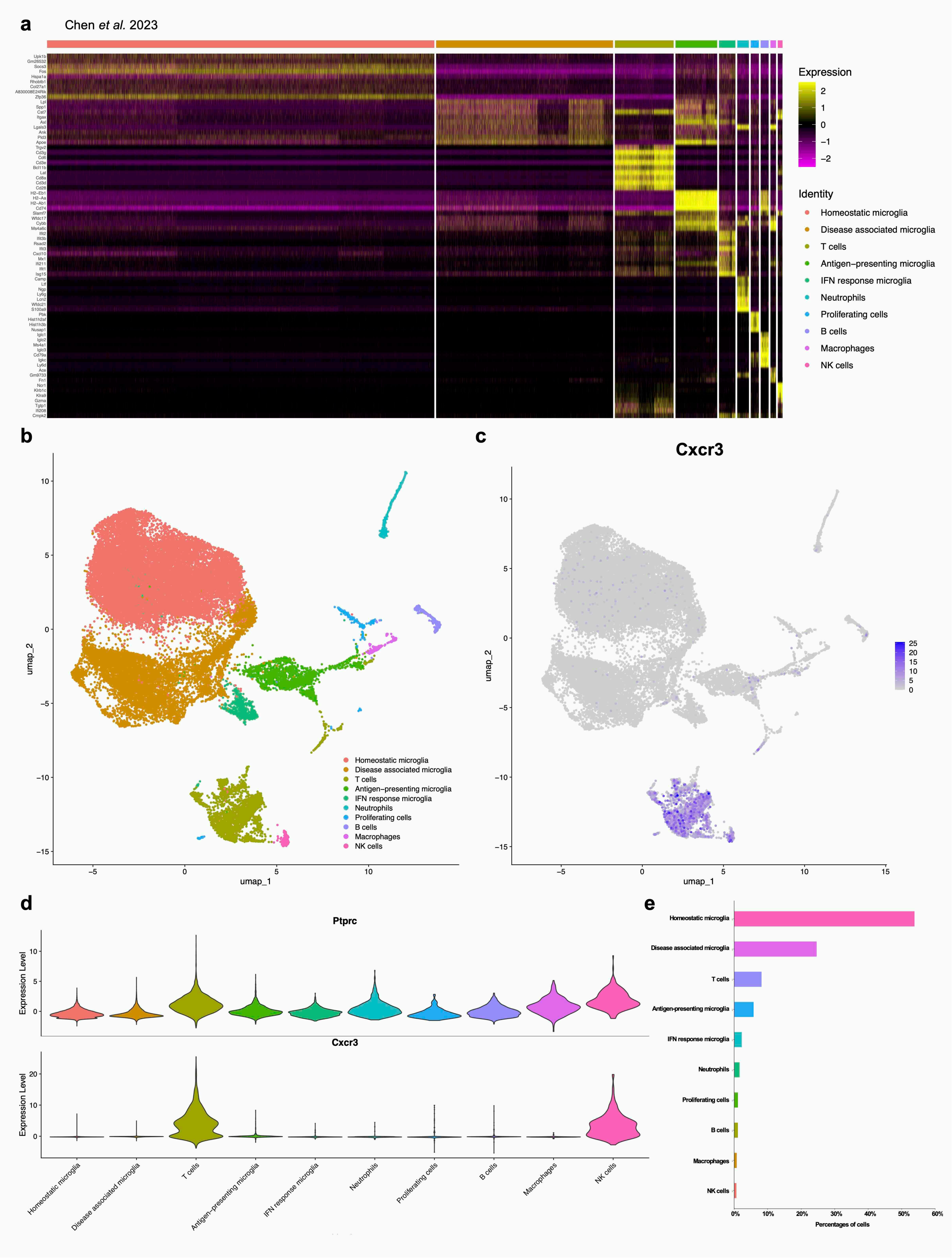

a
Chen et al. 2023
Expression
Identity
Homeostatic microglia
Disease associated microglia
T cells
Antigen−presenting microglia
IFN response microglia
Neutrophils
Proliferating cells
B cells
Macrophages
NK cells
b
umap_1
umap_2
c
Cxcr3
d
Ptprc
Cxcr3
Expression Level
e
Percentages of cells

**Supplementary Fig. 13 | Re-analysis of scRNA-seq datasets to identify *Cxcr3*-expressing cell types in tauopathy mouse model.** **a–d** Re-analysis of the single-cell RNA-seq (scRNA-seq) dataset (GSE221856) from CD45$^+$-sorted cells in TE4 and E4 mice. **a** The top 10 genes upregulated in each cluster are shown in heatmap. **b** Immune cells categorized as CD45$^+$ from TE4 and E4 mice were classified into eight distinct cell types, as illustrated by UMAP plots. These eight cell types were identified based on unique gene expression patterns. **c** Representative mRNA expression patterns of *Cxcr3* in TE4 and E4 mice by Feature plot. **d** Violin plot illustrating cell clusters expressing *Ptprc* and *Cxcr3*. **e** Bar graph showing the percentage of each cell type in TE4 and E4 mice. Number of mice used: male E4 (n = 2), male TE4 (n = 2). Source data are provided in the Source Data file.

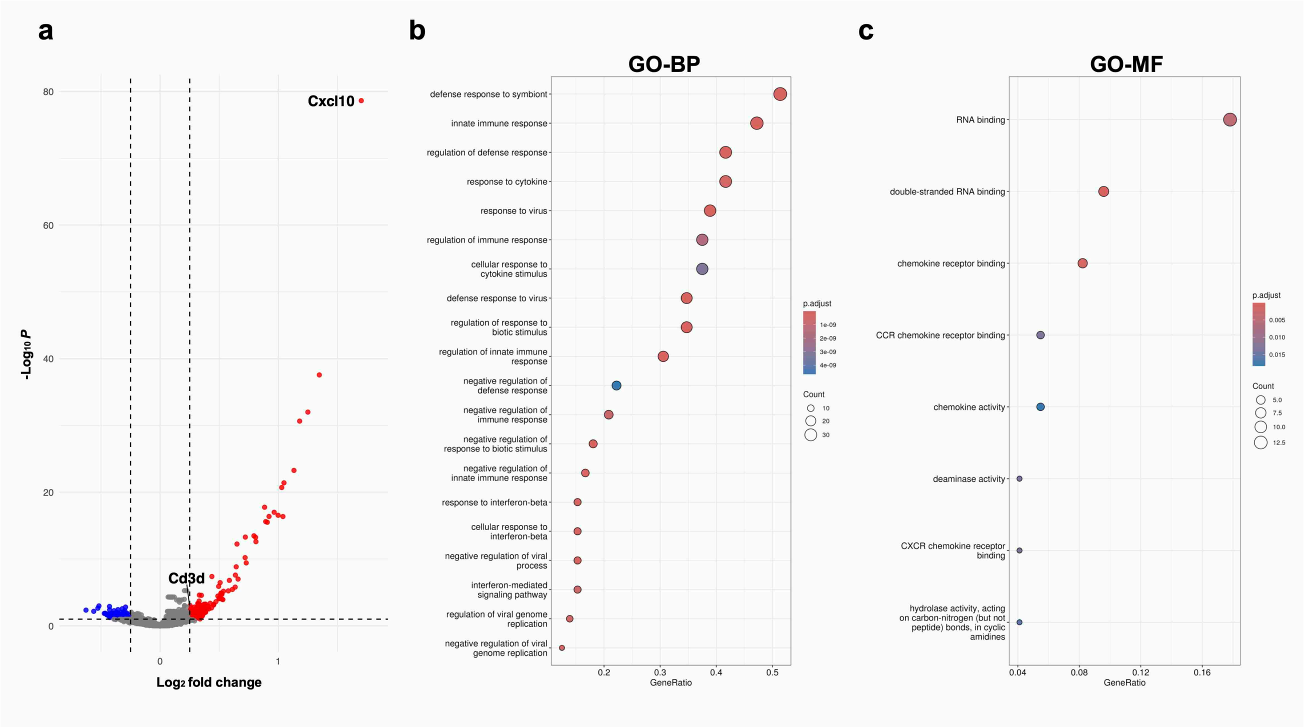


**Supplementary Fig. 14 | Characterization of cells neighboring *Cxcl10*⁺ astrocytes in female Tau Tg mice.**

**a** Volcano plot illustrates $\text{Log}_2$ fold change > |0.25| and p < 0.1 highlighted in red, $\text{Log}_2$ fold change < |0.25| and p > 0.1 shown in blue. All other transcripts are indicated in gray. **b, c** GO pathway analysis focusing on BP and MF was performed to compare cells neighboring *Cxcl10*$^+$ astrocytes with those neighboring *Cxcl10*$^-$ astrocytes in female Tau Tg mice. The top 20 significantly upregulated pathways in cells neighboring *Cxcl10*$^+$ astrocytes are shown in each panel. Number of mice used: female Tau Tg (n = 1). **a-c** DEGs were identified using the Wilcoxon test, and GO analysis was evaluated via over-representation analysis employing the hypergeometric test. Source data are provided in the Source Data file.

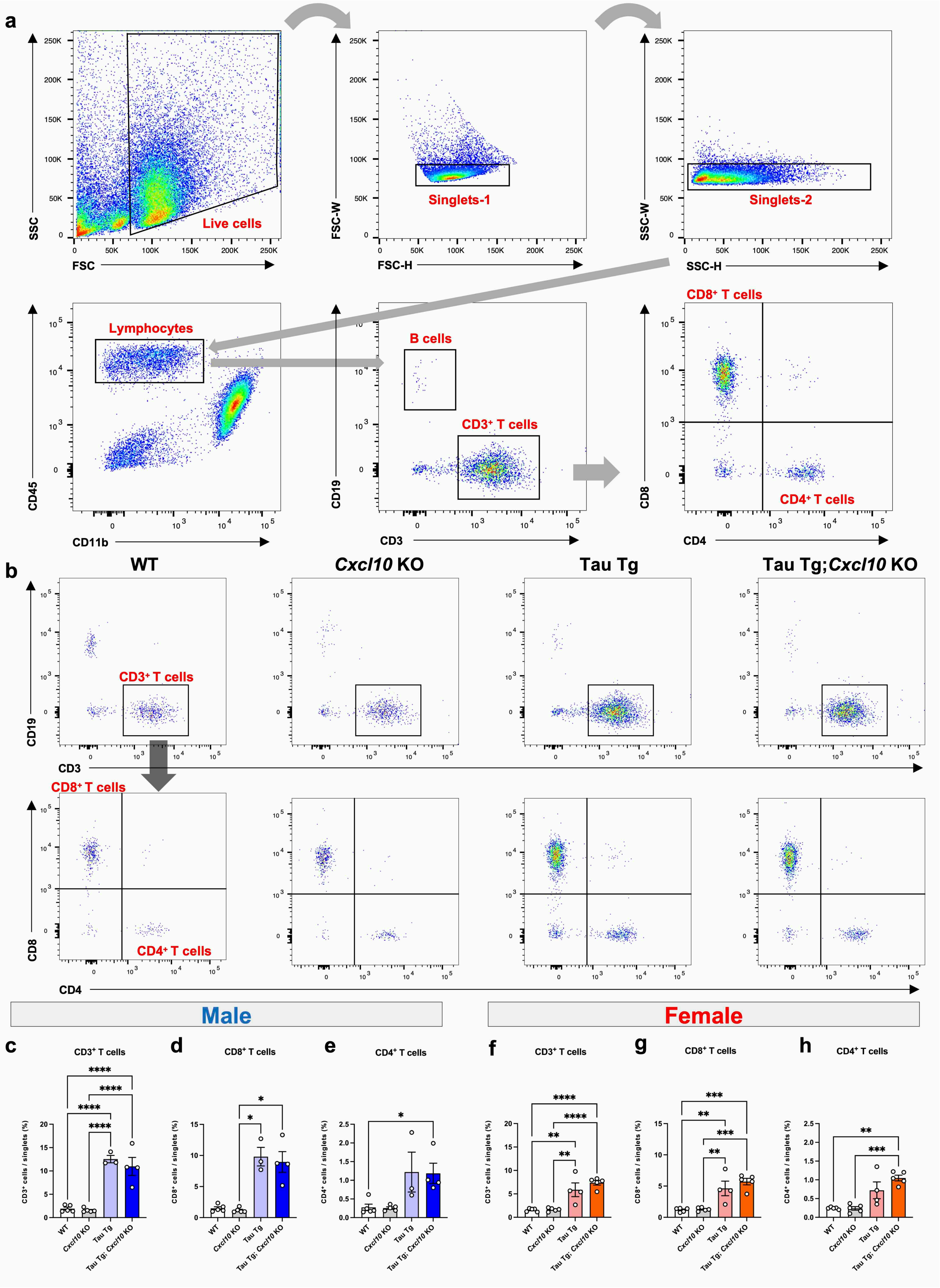

a
Live cells
Singlets-1
Singlets-2
SSC
FSC
FSC-W
FSC-H
SSC-W
SSC-H
Lymphocytes
B cells
CD3+ T cells
CD8+ T cells
CD4+ T cells
CD45
CD11b
CD19
CD3
CD8
CD4
b
WT
Cxcl10 KO
Tau Tg
Tau Tg;Cxcl10 KO
Male
Female
c
CD3+ T cells
d
CD8+ T cells
e
CD4+ T cells
f
CD3+ T cells
g
CD8+ T cells
h
CD4+ T cells
CD3+ cells / singlets (%)
CD8+ cells / singlets (%)
CD4+ cells / singlets (%)
WT
Cxcl10 KO
Tau Tg
Tau Tg; Cxcl10 KO

**Supplementary Fig. 15 | Flow cytometric analysis of T cells in the brain.**

**a** Sequential gating was performed to identify live cells and singlets, followed by gating for $CD45^+$ $CD11b^-$ cells. $CD19^+$ B cells and $CD3^+$ T cells were further classified. Within the $CD3^+$ cell population, $CD4^+$ and $CD8^+$ T cells were identified. **b** Representative gating plots of $CD3^+$, $CD4^+$, and $CD8^+$ T cells in the brains of mice by from each genotype. **c-h** Quantification of single-cell populations: $CD3^+$ T cells (**c**: male, **f**: female), $CD8^+$ T cells (**d**: male, **g**: female), and $CD4^+$ T cells (**e**: male, **h**: female). Data are presented as mean ± S.E.M. Number of mice used: male WT (n = 5), male *Cxcl10* KO (n = 5), male Tau Tg (n = 3), and male Tau Tg; *Cxcl10* KO (n = 4). Female WT (n = 5), female *Cxcl10* KO (n = 5), female Tau Tg (n = 4), and female Tau Tg; *Cxcl10* KO (n = 5). Statistical analyses were performed using a one-way ANOVA followed by Tukey's multiple comparisons test (**b, e, f**), a Brown-Forsythe ANOVA test followed by Dunnett's T3 multiple comparisons test (**g**), and a Kruskal-Wallis test followed by Dunn's multiple comparisons test (**c, d**). $^{**}p < 0.01$, $^{***}p < 0.001$, $^{****}p < 0.0001$. Source data are provided in the Source Data file.

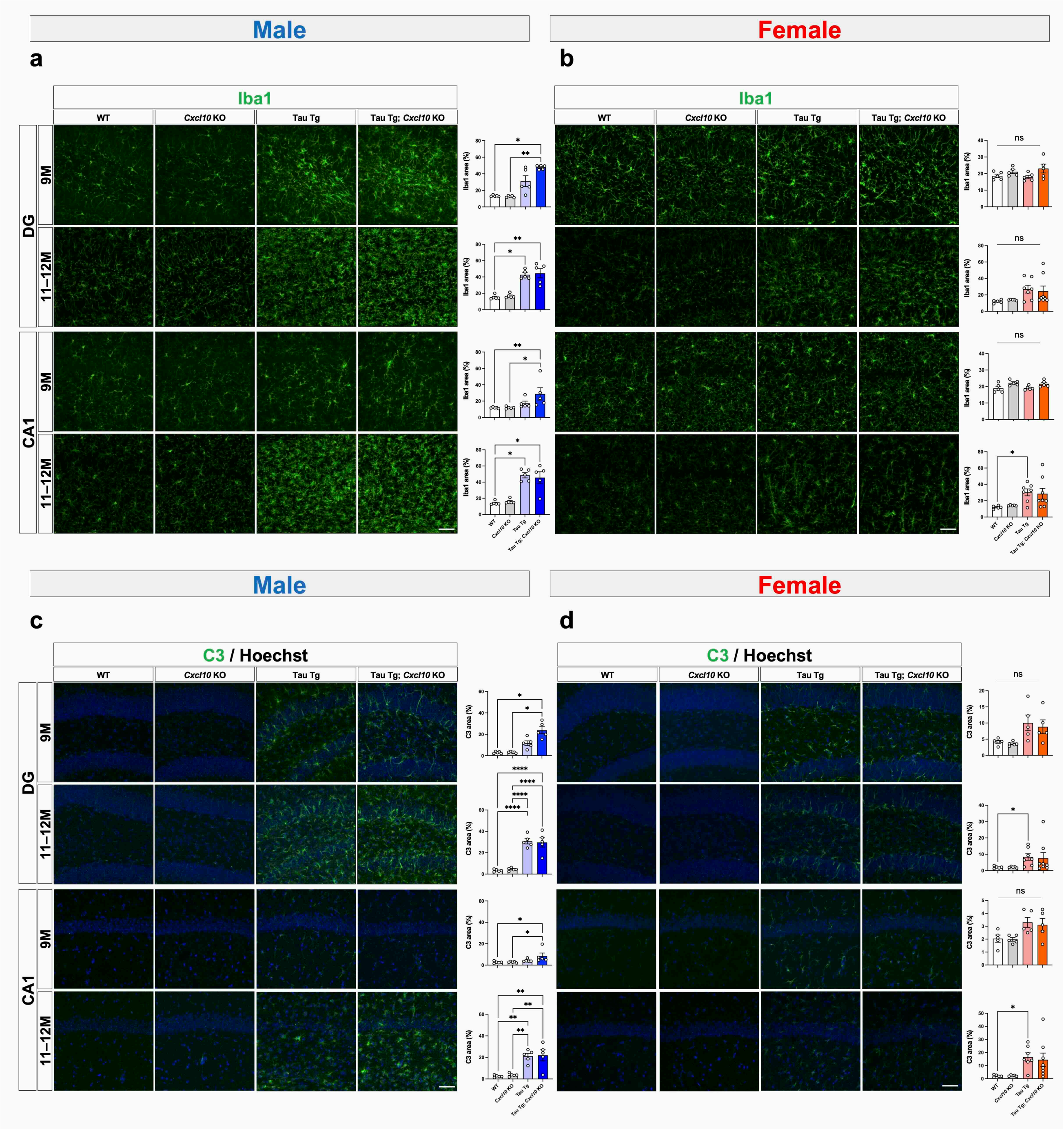


**Supplementary Fig. 16 | Immunohistochemical analysis of microglia and astrocytes in 9- and 11–12-month-old mice.**

**a–d** Representative images of brain sections stained with anti-Iba1 (**a**, **b**) and anti-C3 (**c**, **d**) in the DG and CA1 regions of 9- or 11–12-month-old mice (**a, c** male; **b, d** female). Scale bar = 500 μm. Bar graphs represent the quantified percentage of the image area that is positive for Iba1 or C3. Data are presented as mean ± S.E.M. Number of mice used: male WT (n = 5), male *Cxcl10* KO (n = 5), male Tau Tg (n = 5), and male Tau Tg; *Cxcl10* KO (n = 5). Female WT (n = 5), female *Cxcl10* KO (n = 5), female Tau Tg (n = 7), and female Tau Tg; *Cxcl10* KO (n = 8). Statistical analyses were performed using a one-way ANOVA followed by Tukey's multiple comparisons test (**b**-9M-DG**, b**-9M-CA1**, c**-11–12M-DG**, c**-11–12M-CA1**, d**-9M-DG**, d**-9M-CA1), a Brown-Forsythe ANOVA test followed by Dunnett's T3 multiple comparisons test (**c**-9M-DG), and a Kruskal-Wallis test followed by Dunn's multiple comparisons test (**a, b**-11–12M-DG**, b**-11–12M-CA1**, c**-9M-CA1**, d**-11–12M-DG**, d**-11–12M-CA1). *$p < 0.05$, *$p < 0.01$, ****$p < 0.0001$. Source data are provided in the Source Data file.

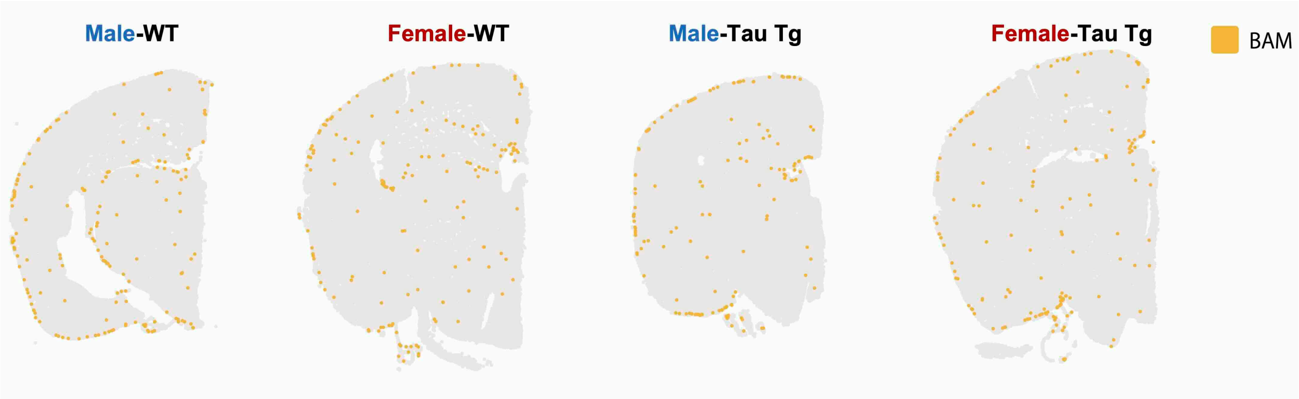


**Supplementary Fig. 17 | Investigating the localization of BAM in mouse brain tissue**

Images illustrating the localization of BAM in brain sections used for the Xenium analysis. Number of mice used: male WT (n = 1), male Tau Tg (n = 1), female WT (n = 1), female Tau Tg (n = 1).

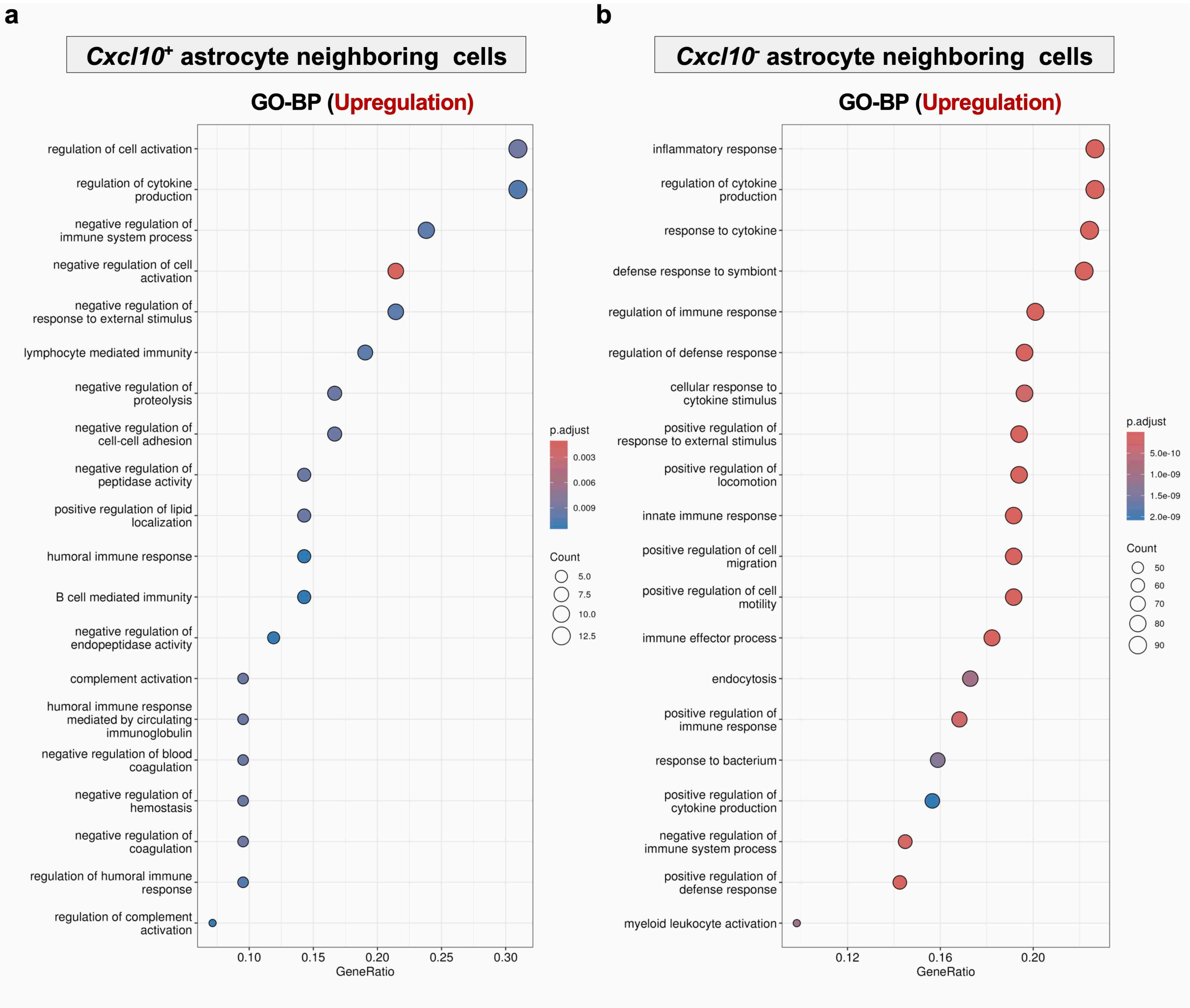


**Supplementary Fig. 18 | Pathway analysis of sex-dependent differences in gene expression in cells neighboring *Cxcl10*⁺ or *Cxcl10*⁻ astrocytes in Tau Tg mice.**

**a**, **b** Genes expressed in cells neighboring *Cxcl10*⁺ astrocytes in females were used as the baseline to identify differential expression in the corresponding male condition, followed by BP pathway of GO analysis (**a**). The same analytical framework was applied to *Cxcl10*⁻ astrocytes (**b**). The top 20 significantly upregulated pathways in cells neighboring *Cxcl10*⁺ astrocytes (**a**) and *Cxcl10*⁻ astrocytes (**b**) in males are shown in each panel. Number of mice used: male Tau Tg (n = 1), female Tau Tg (n = 1). **a**, **b** DEGs were identified using the Wilcoxon test, and GO analysis was evaluated via over-representation analysis employing the hypergeometric test. Source data are provided in the Source Data file.

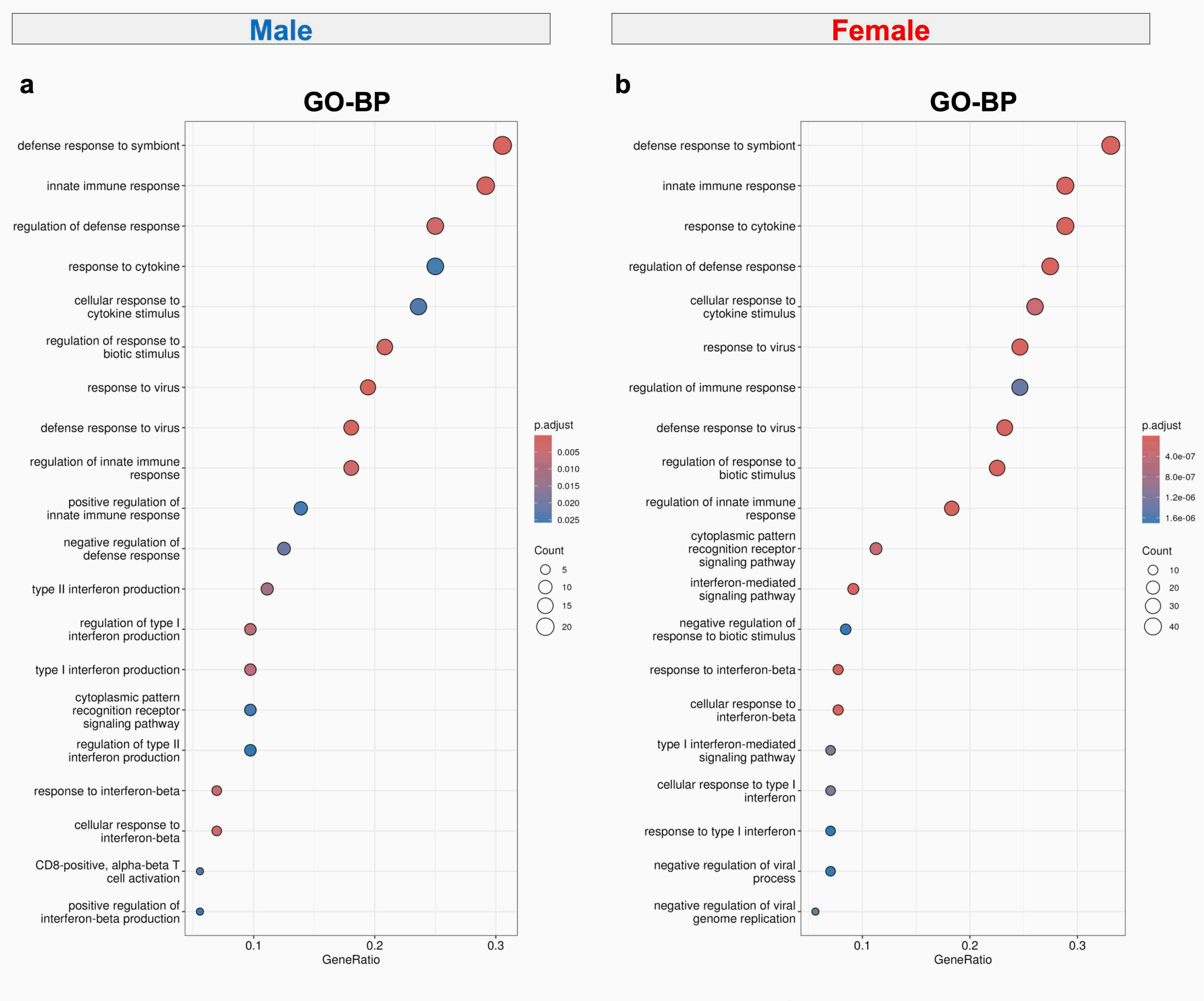


**Supplementary Fig. 19 | Pathway analysis to investigate characterization of *Cxcl10*⁺ astrocytes in the brains of Tau Tg mice.**

**a** GO pathway analysis focusing on BP was performed to compare *Cxcl10*$^{+}$ astrocytes with *Cxcl10*$^{-}$ astrocytes in Tau Tg mice. The top 20 significantly upregulated pathways in male (**a**) and female (**b**) mice are shown in each panel. Number of mice used: male Tau Tg (n = 1), female Tau Tg (n = 1). **a**,**b** DEGs were identified using the Wilcoxon test, and GO analysis was evaluated via over-representation analysis employing the hypergeometric test. Source data are provided in the Source Data file.

### Supplementary Tables

##### Supplementary Table 1 | List of antibodies used for western blotting

| Antibodies | Source | Dilution | Identifier |
|---|---|---|---|
| anti-human Phospho-Tau (Ser202, Thr205) IgG [AT8] | FUJIREBIO, Inc. | 1: 1,000 | Cat# 90206, RRID: AB_223648 |
| anti-human/mouse/rat phospho-Tau (Ser396) IgG | Thermo Fisher Scientific, Inc. | 1: 5,000 | Cat# 44-752G, RRID:AB_2533745 |
| anti-human/mouse/rat Tau IgG [TAU-5] | Abcam, plc. | 1: 1,000 | Cat# ab80579, RRID:AB_1603723 |
| anti-human Phospho-Tau (Thr212, Ser214) IgG [AT100] | Thermo Fisher Scientific, Inc. | 1: 1,000 | Cat# MN1060, RRID:AB_223652 |
| anti-human/mouse/rat Phospho-Tau (Thr181) IgG [D9F4G] | Cell Signaling Technology, Inc. | 1: 1,000 | Cat# 12885, RRID:AB_2798053 |
| anti-mouse IgG (H+L), Horseradish Peroxidase | Jackson ImmunoResearch, Inc. | 1: 20,000 | Cat# 715-035-150, RRID:AB_2340770 |
| anti-rabbit IgG (H+L), Horseradish Peroxidase | Jackson ImmunoResearch, Inc. | 1: 20,000 | Cat# 711-035-152, RRID:AB_10015282 |

**Supplementary Table 2 | List of antibodies used for immunohistochemical analysis**

| Antibodies [clone] | Source | Dilution | Identifier |
|---|---|---|---|
| anti-human CD3 IgG [CD3-12] | Bio-Rad Laboratories, Inc. | 1: 500 | Cat# MCA1477T, RRID:AB_10845948 |
| anti-mouse/rat CD31/PECAM1 IgG | R&D Systems, Inc. | 1: 500 | Cat# AF3628, RRID:AB_2161028 |
| anti-human/mouse/rat NeuN IgG [D4G4O] | Cell Signaling Technology, Inc. | 1: 500 | Cat# 24307, RRID:AB_2651140 |
| anti-human/rat Iba1 IgG | Abcam, plc. | 1: 1,000 | Cat# ab5076, RRID:AB_2224402 |
| Anti Iba1, Rabbit (for Immunocytochemistry) | FUJIFILM Wako Pure Chemical, Corp. | 1: 500 | Cat# 019-19741, RRID:AB_839504 |
| anti-GFAP antibody :: Goat GFAP Polyclonal Antibody | MyBioSource, Inc. | 1: 1,000 | Cat# MBS241916, RRID:AB_839504 |
| anti-mouse C3 IgG [11H9] | Hycult Biotech, Inc. | 1: 1,000 | Cat# HM1045, RRID:NA |
| anti-human Phospho-Tau (Ser202, Thr205) IgG [AT8], biotin | FUJIREBIO, Inc. | 1: 200 | Cat# 90206, RRID: AB_223648 |
| anti-rat IgG (H+L) Highly Cross-Adsorbed Secondary Antibody, Alexa Fluor™ 488 | Thermo Fisher Scientific, Inc. | 1: 1,000 | Cat# A-21208, RRID:AB_2535794 |
| anti-rabbit IgG (H+L) Highly Cross-Adsorbed Secondary Antibody, Alexa Fluor™ 488 | Thermo Fisher Scientific, Inc. | 1: 1,000 | Cat# A-21206, RRID:AB_2535792 |
| anti-rabbit IgG (H+L) Cross-Adsorbed Secondary Antibody, Alexa Fluor™ 568 | Thermo Fisher Scientific, Inc. | 1: 1,000 | Cat# A-10042, RRID:AB_2534017 |
| anti-goat IgG (H+L) Highly Cross-Adsorbed Secondary Antibody, Alexa Fluor™ 488 | Thermo Fisher Scientific, Inc. | 1: 1,000 | Cat# A-11055, RRID:AB_2534102 |
| anti-goat IgG (H+L) Cross-Adsorbed Secondary Antibody, Alexa Fluor™ 568 | Thermo Fisher Scientific, Inc. | 1: 1,000 | Cat# A-11057, RRID:AB_2534104 |
| anti-goat IgG (H+L) Cross-Adsorbed Secondary Antibody, Alexa Fluor™ 647 | Thermo Fisher Scientific, Inc. | 1: 1,000 | Cat# A-21447, RRID:AB_2535864 |

**Supplementary Table 3 | List of antibodies used for flow cytometry**

| Antibodies | Source | Dilution | Identifier |
|---|---|---|---|
| anti-mouse CD45 IgG [30-F11], FITC | BioLegend, Inc. | 1: 400 | Cat# 11-0451-82, RRID:AB_465050 |
| anti-mouse/human CD11b IgG [M1/70], PE/Cyanine7 | BioLegend, Inc. | 1: 800 | Cat# 101216, RRID:AB_312799 |
| anti-mouse CD19 IgG [6D5], Brilliant Violet 785™ | BioLegend, Inc. | 1: 100 | Cat# 115543, RRID:AB_11218994 |
| anti-human/mouse CD3e IgG [145-2C11], APC | Thermo Fisher Scientific, Inc. | 1: 400 | Cat# 17-0031-82, RRID:AB_469315 |
| anti-mouse CD8a IgG [53-6.7], Brilliant Violet 510™ | Becton, Dickinson, Co. | 1: 100 | Cat# 563068, RRID:AB_2687548 |
| anti-mouse CD4 IgG [RM4–5], APC/Cyanine7 | BioLegend, Inc. | 1: 400 | Cat# 100525, RRID:AB_312726 |

**Supplementary Table 4 | Title: List of materials materials used**

| **Resource or laboratory equipment** | **Source** | **Identifier** |
|---|---|---|
| **Mouse** | | |
| C57BL/6J | The Jackson Laboratory Japan, Inc | Cat# 000664, RRID:IMSR_JAX:000664 |
| B6.129S4-*Cxcl10*tm1Adl/J | The Jackson Laboratory Japan, Inc | Cat# 006087, RRID:IMSR_JAX:006087 |
| B6.129X1-Mapttm1Hnd/J | The Jackson Laboratory Japan, Inc | Cat# 007251, RRID:IMSR_JAX:007251 |
| B6.Cg-Tg(Prnp-MAPT*P301S)PS19Vle/J | Dr. Virginia M Y Lee | Cat# 024841, RRID:IMSR_JAX:024841 |
| **Software** | | |
| GraphPad Prism (version 10.4.0) | GraphPad Software, Llc | Cat# NA, RRID:SCR_002798 |
| Image Lab software | Bio-Rad Laboratories, Inc. | Cat# 12012931, RRID:SCR_014210 |
| FlowJo | Becton, Dickinson, Co. | Cat# 663335, RRID:SCR_008520 |
| Smart3.0 Video Tracking Software | Harvard Apparatus \| Panlab, S.L.U. | Cat# SMARTBASIC, RRID:NA |
| R (version:4.5.2) | | Cat# NA, RRID:NA |
| NDP.view | Hamamatsu Photonics, K.K. | Cat# NA, RRID:NA |
| Fiji | | Cat# NA, RRID:SCR_002285 |
| Loupe Browser 9.0 | 10x Genomics, Inc. | Cat# NA,RRID: SCR_018555 |
| Xenium Explorer 4.1.1 | 10x Genomics, Inc. | Cat# NA, RRID: SCR_025847 |
| **Equipment** | | |
| Multi-beads shocker | Yasui Kikai, Corp. | Cat# MB3000, RRID:NA |
| Ultracentrifuge | Hitachi, Ltd. | Cat# CP100WX, RRID:NA |
| ChemiDoc™ Touch MP imaging system | Bio-Rad Laboratories, Inc. | Cat# 17001402JA, RRID:SCR_021693 |
| Cryostat | Leica Microsystems, Inc. | Cat# CM1850, RRID:NA |
| FV3000 confocal laser scanning microscope | Evident Scientific, Inc. | Cat# 7M88501, RRID:SCR_017015 |
| Virtual slide scanner NanoZoomer S60 | Hamamatsu Photonics, K.K. | Cat# C13210-01, RRID:SCR_023762 |
| GRIP STRENGTH METER FOR RATS & MICE MODEL | Muromachi Kikai Co., Ltd. | Cat# MK-380V, RRID:NA |
| ROTA-ROD TREADMILL FOR MICE | Muromachi Kikai Co., Ltd. | Cat# MK-610A, RRID:NA |
| NanoDrop One | Thermo Fisher Scientific, Inc. | Cat# ND-ONE-W, RRID:SCR_023005 |
| FACSAria III Cell Sorter | Becton, Dickinson, Co. | Cat# 648282, RRID:SCR_016695 |
